# Somatosensory-Motor Dysconnectivity Spans Multiple Transdiagnostic Dimensions of Psychopathology

**DOI:** 10.1101/637827

**Authors:** Valeria Kebets, Avram J. Holmes, Csaba Orban, Siyi Tang, Jingwei Li, Nanbo Sun, Ru Kong, Russell A. Poldrack, B.T. Thomas Yeo

**Affiliations:** Department of Electrical and Computer Engineering, ASTAR-NUS Clinical Imaging Research Centre, Singapore Institute for Neurotechnology and Memory Networks Program, National University of Singapore, Singapore; Department of Psychology, Yale University, New Haven, CT, USA; Department of Psychiatry, Yale University, New Haven, CT, USA; Athinoula A. Martinos Center for Biomedical Imaging, Massachusetts General Hospital, Charlestown, MA, USA; Department of Psychiatry, Massachusetts General Hospital, Boston, MA, USA; Department of Psychology, Stanford University, CA, USA; Centre for Cognitive Neuroscience, Duke-NUS Medical School, Singapore; NUS Graduate School for Integrative Sciences and Engineering, National University of Singapore, Singapore

**Keywords:** Psychopathology, Cognitive dysfunction, Impulsivity, Somatosensory-motor, Phenotypes, Resting state functional connectivity

## Abstract

**Background:** There is considerable interest in a dimensional transdiagnostic approach to psychiatry. Most transdiagnostic studies have derived factors based only on clinical symptoms, which might miss possible links between psychopathology, cognitive processes and personality traits. Furthermore, many psychiatric studies focus on higher-order association brain networks, thus neglecting the potential influence of huge swaths of the brain.

**Methods:** A multivariate data-driven approach (partial least squares; PLS) was utilized to identify latent components linking a large set of clinical, cognitive and personality measures to whole-brain resting-state functional connectivity (RSFC) patterns across 224 participants. The participants were either healthy (N=110) or diagnosed with bipolar disorder (N=40), attention-deficit/hyperactivity disorder (N=37), schizophrenia (N=29) or schizoaffective disorder (N=8). In contrast to traditional case-control analyses, the diagnostic categories were *not* utilized in the PLS analysis, but were helpful for interpreting the components.

**Results:** Our analyses revealed three latent components corresponding to general psychopathology, cognitive dysfunction and impulsivity. Each component was associated with a unique whole-brain RSFC signature and shared across all participants. The components were robust across multiple control analyses and replicated using independent task functional magnetic resonance imaging data from the same participants. Strikingly, all three components featured connectivity alterations within the somatosensory-motor network, and its connectivity with subcortical structures and cortical executive networks.

**Conclusions:** We identified three distinct dimensions with dissociable (but overlapping) whole-brain RSFC signatures across healthy individuals and individuals with psychiatric illness, providing potential intermediate phenotypes that span across diagnostic categories. Our results suggest expanding the focus of psychiatric neuroscience beyond higher-order brain networks.

## Introduction

Substantial overlap in clinical symptoms (1,2), cognitive deficits (3), and genetic risk factors (4) across psychiatric disorders, as well as the high comorbidity rates (5) suggest that current categorical classifications might not be carving nature by its joints. In response, transdiagnostic initiatives, such as the Research Domain Criteria (6,7) and the Hierarchical Taxonomy of Psychopathology (8,9), have worked towards new dimensionally-oriented approaches that would integrate findings from genetics, neuroimaging and cognitive science.

Many recent transdiagnostic studies have derived latent dimensional factors that best explain the structure of psychopathology, along with associated neural correlates. One example is the general psychopathology (or *p*) factor (10–15), which is thought to reflect individuals’ susceptibility to develop “any and all forms of common psychopathologies” (16). The *p* factor has also been extended to other clinical dimensions, e.g., internalizing/ externalizing symptoms (10,11,17). Importantly, these factors were extracted from the general population, supporting the idea that psychopathology lies on a spectrum spanning healthy and disease states. However, most studies have focused on deriving factors based only on clinical symptoms, which might be insensitive to intricate links between psychopathology, cognitive processes and personality traits. Therefore, considering a broader set of behavioral measures might provide a more comprehensive characterization of individuals’ phenotypic variability across mental health and disease.

Resting-state functional magnetic resonance imaging (rs-fMRI) is widely used to measure intrinsically organized patterns of spontaneous signal fluctuations (18), commonly referred to as resting-state functional connectivity (RSFC). Since many psychiatric disorders are characterized by disturbances of large-scale brain network organization (19), RSFC might be a powerful tool for understanding transdiagnostic dimensions. Indeed, studies have found significant overlap in neural circuits altered in different disorders, suggesting common neurobiological mechanisms (20–23). RSFC alterations in higher-order (e.g., default mode and executive) networks are also associated with the *p* factor (14,17) or clinical symptoms (24). However, many psychiatric imaging studies have focused on higher-order association networks (19,25), neglecting the potential influence of huge swaths of cortex. Indeed, complex clinical and behavioral phenotypes arise from coordinated interactions throughout the entire connectome (26–28), suggesting the importance of examining whole-brain connectivity without prior assumptions.

In this study, we utilized data from the UCLA Consortium for Neuropsychiatric Phenomics (29), a unique dataset in which the imaging and behavioral assessment were focused on working memory and inhibitory control, two domains that are exceedingly relevant in multiple psychiatric disorders. This allowed us to examine behavioral phenotypes beyond the clinical symptoms typically examined in many transdiagnostic studies (10,11,14,15,17). Using a multivariate data-driven approach, we extracted *latent components* that simultaneously link a large set of behavioral measures spanning clinical, cognitive and personality domains with whole-brain RSFC patterns across healthy individuals (HC) and individuals with schizophrenia (SZ), schizoaffective disorder (SZAD), bipolar I disorder (BD) or attention deficit/hyperactivity disorder (ADHD). In contrast to traditional case-control analyses, the diagnostic labels were *not* utilized in the analysis, but used to interpret the behavioral-RSFC dimensions posthoc.

Our analyses revealed three transdiagnostic components corresponding to general psychopathology, cognitive dysfunction and impulsivity. Each component was associated with a unique whole-brain RSFC pattern, such that inter-individual variation in the expression of the three RSFC configurations captured individuals’ variability along these behavioral dimensions. Strikingly, the three components all featured altered connectivity within the somatosensory-motor system, and in its connections to subcortical and cortical executive networks. Overall, this study identifies three latent components as likely transdiagnostic phenotypes, thereby offering a putative model for explaining comorbidity across disorders. Our results add further evidence to the importance of considering a broad range of behavioral measures, as well as expanding the focus of psychiatric neuroscience beyond higher-order association brain networks.

## Methods and Materials

### Participants

We downloaded the UCLA Consortium for Neuropsychiatric Phenomics (CNP) dataset from the public database OpenfMRI (30). The CNP dataset (29) comprised multimodal imaging and behavioral data from 272 participants, including 130 healthy individuals (HC), 49 patients with bipolar disorder type I (BD), 43 patients with attention-deficit/hyperactivity disorder (ADHD), 39 patients with schizophrenia (SZ) and 11 patients with schizoaffective disorder (SZAD).

Details about participant recruitment can be found elsewhere (29). Briefly, healthy adults were recruited via community advertisements from the Los Angeles area, while patients were recruited via local clinics and online portals. Inclusion criteria were: age between 21-50 years old; NIH racial/ethnic category either White (not Hispanic or Latino), or Hispanic or Latino (of any racial group); primary language either English or Spanish; completed at least 8 years of formal education; no significant medical illness. Participants were screened for drugs of abuse (cannabis, amphetamine, opioids, cocaine, benzodiazepines), and excluded in cases where urinalysis results were positive. Other exclusion criteria were: being left-handed, pregnancy, history of head injury with loss of consciousness or cognitive sequelae, or other MRI contraindications (e.g., claustrophobia). Stable medications were permitted for patients.

After receiving a verbal explanation of the study, participants gave written informed consent following procedures approved by the Institutional Review Boards at UCLA and the Los Angeles County Department of Mental Health.

### Clinical and behavioral assessment

All participants underwent a semi-structured assessment with the Structured Clinical Interview for the Diagnostic and Statistical Manual of Mental Disorders, Fourth Edition – Text Revision (SCID-I (31)). Demographic and clinical data for each group are summarized in **Table 1**.

**Table 1.** Demographic and Clinical Data for Each Diagnostic Group.

| | HC<br>(n=110) | ADHD<br>(n=37) | BD<br>(n=40) | SZ<br>(n=29) | SZAD<br>(n=8) | *F* / $\chi^2$ | *p* value |
| --- | --- | --- | --- | --- | --- | --- | --- |
| *Demographics* |  |  |  |  |  |  |  |
| Age, mean (SD), yrs | 31.26 (8.66) | 31.22 (10.11) | 34.78 (9.24) | 35.59 (9.27) | 35.38 (8.94) | 2.34 | 5.6E-02 |
| Female sex, No. (%) | 53 (48%) | 17 (46%) | 17 (43%) | 4 (14%) | 4 (50%) | 11.60 | 2.1E-02 |
| Education, mean (SD), yrs | 15.19 (1.59) | 14.57 (1.85) | 14.43 (1.91) | 12.62 (1.47) | 13.38 (1.77) | 14.49 | **1.6E-10** |
| Site 1, No. (%) | 89 (81%) | 19 (51%) | 18 (45%) | 13 (45%) | 4 (50%) | 27.74 | **1.4E-05** |
| *Head motion* |  |  |  |  |  |  |  |
| FD, mean (SD) <sup>a</sup> | 0.06 (0.03) | 0.06 (0.03) | 0.07 (0.03) | 0.08 (0.03) | 0.09 (0.05) | 3.50 | **8.5E-03** |
| *Lifetime substance use* <sup>b</sup> |  |  |  |  |  |  |  |
| 0 substance, No. (%) | 69 (63%) | 15 (41%) | 6 (15%) | 9 (31%) | 2 (25%) | 32.38 | **1.6E-06** |
| 1 substance, No. (%) | 24 (22%) | 11 (30%) | 8 (20%) | 5 (17%) | 0 (0%) | 4.06 | 4.0E-01 |
| 2+ substances, No. (%) | 17 (15%) | 11 (30%) | 26 (65%) | 15 (52%) | 6 (75%) | 44.65 | **4.7E-09** |
| # substances, mean (SD) | 0.62 (1.04) | 1.32 (1.63) | 2.73 (2.11) | 2.07 (2.15) | 3.00 (2.51) | 16.67 | **6.1E-12** |
| *Current medication (by target)* <sup>c</sup> |  |  |  |  |  |  |  |
| Dopaminergic, No. (%) | 0 (0%) | 10 (27%) | 17 (42%) | 19 (66%) | 6 (75%) | 44.80 | **1.9E-27** |
| Serotonergic, No. (%) | 0 (0%) | 2 (5%) | 14 (35%) | 17 (59%) | 2 (25%) | 71.14 | **1.6E-38** |
| GABAergic, No. (%) | 0 (0%) | 2 (5%) | 13 (32%) | 5 (17%) | 0 (0%) | 13.50 | **7.4E-10** |
| Glutamatergic, No. (%) | 0 (0%) | 1 (3%) | 13 (32%) | 3 (10%) | 2 (25%) | 25.11 | **3.9E-17** |
| Norepinephrinergic, No. (%) | 0 (0%) | 10 (27%) | 14 (35%) | 9 (31%) | 4 (50%) | 16.64 | **6.4E-12** |
| Others, No. (%) | 0 (0%) | 1 (3%) | 5 (12%) | 5 (17%) | 1 (12%) | 4.87 | **8.8E-04** |
| # meds, mean (SD) | 0 (0) | 0.59 (1.14) | 2.33 (1.85) | 2.00 (1.51) | 2.50 (1.77) | 47.19 | **1.4E-28** |

*Abbreviations:* HC, healthy controls; ADHD, attention deficit/hyperactivity disorder; BD, bipolar disorder; SZ, schizophrenia; SZAD, schizoaffective disorder; SD, standard deviation; yrs, years; FD, framewise displacement; #, number of; meds, medications.
Groups were compared with either ANOVAs (for continuous measures) or chi-squared tests (for categorical measures). All *p*-values that survived FDR correction ( $q < 0.05$ ) are indicated in bold.
<sup>a</sup> Framewise displacement was computed as per Kong et al. (32).
<sup>b</sup> Lifetime substance use included substance abuse and/or dependence for nicotine, alcohol, cannabis, cocaine, amphetamine, sedatives/hypnotics/anxiolytics, inhalants, opioids and hallucinogens.
<sup>c</sup> Medication was sorted by the neurotransmitter system(s) targeted by the medication currently used by participants, based on the Neuroscience-based Nomenclature (NbN-2 (33,34); http://www.nbn2.com/). The full list of medications and their categorization can be found in **Table S3**. Note that percentages do not add up to 100% because individuals often take more than one medication.

The CNP behavioral assessment included an extensive set of clinical, cognitive, and personality scores (listed in **Table S1**). We excluded behavioral measures from the partial least square (PLS) analysis when scores were missing for at least one participant among the 224 participants who survived MRI preprocessing quality controls (see below). Notably, most clinical measures were excluded from the PLS analysis as they had not been administered to healthy individuals.

A final set of 54 behavioral and self-report measures from 19 clinical, cognitive and psychological tests were included in the PLS analysis. Behavioral measures for each group are shown in **Table S2.**

### MRI data acquisition

Structural and functional MRI data were acquired on two 3T Siemens Trio scanners, located at the Ahmanson-Lovelace Brain Mapping Center (Siemens version Syngo MR B15) and the Staglin Center for Cognitive Neuroscience (Siemens version Syngo MR B17) at UCLA, on two separate days, in a counter-balanced fashion. fMRI acquisition comprised a resting-state fMRI (rs-fMRI) scan and 7 fMRI tasks (Balloon Analog Risk, Paired Associate Memory Encoding and Retrieval, Stop Signal, Spatial Capacity Working Memory, Task Switching and Breath Holding). Details about the task paradigms can be found elsewhere (29).

fMRI data were collected using a T2*-weighted echoplanar imaging (EPI) sequence with the following parameters: slice thickness = 4 mm, 34 slices, voxel size = 3 × 3 × 4 mm, TR = 2000 ms, TE = 30 ms, flip angle = 90°, matrix 64 × 64, FOV = 192 mm, oblique slice orientation. The rs-fMRI scan lasted for 304 seconds (152 frames), during which participants were not presented with any stimuli but were asked to keep their eyes open and remain still. A high-resolution MPRAGE anatomical scan was also collected using the following sequence: TR = 1.9 s, TE = 2.26 ms, FOV = 250 mm, matrix = 256 × 256, sagittal plane, slice thickness = 1 mm, 176 slices.

### MRI processing

Of the 272 participants, 7 participants did not have an anatomical scan and 4 were missing a rs-fMRI scan. One participant was also excluded because of signal dropout in the cerebellum (as reported by Gorgolewski et al. (35)). This resulted in 260 participants undergoing preprocessing. The neuroimaging data were processed using a previously published pipeline (32,36) using tools from FSL and FreeSurfer 5.3.0 (http://surfer.nmr.mgh.harvard.edu). The pipeline code is available here: https://github.com/ThomasYeoLab/CBIG/tree/master/stable_projects/preprocessing/CBIG_fMRI_Preproc2016.

Structural MRI data were processed using FreeSurfer 5.3.0, a suite of automated algorithms for reconstructing accurate surface mesh representations of the cortex from individual participants’ T1 images (37–39). The cortical surface meshes were then registered to a common spherical coordinate system (40,41).

Rs-fMRI data were pre-processed with the following steps: (i) removal of first 4 frames, (ii) slice time correction with the FSL package (42,43), (iii) motion correction using rigid body translation and rotation with the FSL package. The structural and functional images were aligned using boundary-based registration (44) using the FsFast software package (http://surfer.nmr.mgh.harvard.edu/fswiki/FsFast). Framewise displacement (FD) and voxel-wise differentiated signal variance (DVARS) were computed using fsl_motion_outliers (43). Volumes with FD > 0.2mm or DVARS > 50 were marked as outliers, as well as one volume before and two volumes after. Uncensored segments of data lasting fewer than 5 contiguous volumes were also flagged as outliers (45). Scans with more than half of the volumes flagged as outliers were removed completely. 36 participants were excluded for excessive head motion, which resulted in a final sample of 224 participants (110 HC, 40 BD, 37 ADHD, 29 SZ, and 8 SZAD patients).

We regressed out nuisance regressors, which consisted of six motion parameters, averaged ventricular signal, averaged white matter signal and global signal, as well as their temporal derivatives (18 regressors in total). The flagged outlier volumes were ignored during the regression procedure. The data were interpolated across censored frames using least squares spectral estimation of the values at censored frames (36). A band-pass filter (0.009 Hz ≤ f ≤ 0.08 Hz) was applied. Finally, the preprocessed fMRI data were projected onto the FreeSurfer fsaverage6 surface space (2mm vertex spacing). Because global signal regression (GSR) is somewhat controversial (46), we also performed control analyses using an alternative strategy (see “Control and reliability analyses”).

### Resting-state functional connectivity (RSFC)

RSFC (Pearson’s correlation) was computed among the average timeseries of 400 cortical (47) and 19 subcortical (48) regions-of-interest (ROIs) covering the entire brain, resulting in a 419 × 419 RSFC matrix for each participant. Because age, sex, education, site, and head motion (mean framewise displacement (42)) were different across groups (**Table 1**), they were regressed from both behavioral and RSFC data. Since the RSFC matrices were symmetric, only the upper triangular portions of the matrices were considered in subsequent analyses (although full matrices are shown for visualization).

### Partial least squares (PLS)

PLS is a multivariate data-driven statistical technique that aims to maximize the covariance between two matrices by deriving *latent components* (LCs), which are optimal linear combinations of the original matrices (49,50). We applied PLS to the RSFC and behavioral measures of all participants *ignoring* diagnostic categories. Each LC is characterized by a distinct RSFC pattern (called RSFC *saliences*) and a distinct behavioral profile (called behavioral *saliences*). By linearly projecting the RSFC and behavioral measures of each participant onto their respective saliences, we obtain individual-specific RSFC and behavioral *composite scores* for each LC. By construction, PLS seeks to find saliences that maximize across-participant covariance between the RSFC and behavioral composite scores. The number of significant LCs was determined by a permutation test accounting for diagnostic categories (1000 permutations). The p-values (from the permutation test) for the top five LCs were corrected for multiple comparisons using a false discovery rate (FDR) of q < 0.05. See **Supplemental Methods** for details.

To interpret the LCs, we computed Pearson’s correlations between the original RSFC data and RSFC composite scores, as well as between the original behavioral measures and behavioral composite scores for each LC (51,52). A large positive (or negative) correlation for a particular behavioral measure for a given LC indicates greater importance of the behavioral measure for the LC. Similarly, a large positive (or negative) correlation for a particular RSFC measure for a given LC indicates greater importance of the RSFC measure for the LC.

### Posthoc analyses

Two-sample t-tests were performed to test if RSFC and behavioral composite scores for LCs 1-3 were different between participants with different diagnoses, medication and substance use. More details can be found in the **Supplemental Methods**.

We also tested if the composite scores were associated with age, sex, years of education, acquisition site, head motion, and medication load. Pearson’s correlations were performed for continuous measures and t-tests for binary measures. As control analyses, we applied GLMs with linear hypothesis tests to assess the impact of diagnosis (while controlling for medication and substance use), medication (while controlling for diagnosis and substance use) and substance use (while controlling for diagnosis and medication). See **Supplemental Methods** for more details.

All posthoc analyses utilized all participants and FDR (q < 0.05) correction was applied to all posthoc tests.

### Control and reliability analyses

Several analyses were performed to ensure robustness of the LCs (see **Supplemental Methods**). First, we tested whether we could replicate the brain-behavior associations identified with rs-fMRI using task-fMRI data. Second, we performed a 5-fold cross-validation of the PLS analysis. Third, we applied quantile normalization to improve the Gaussianity of the behavioral data distributions before PLS. Moreover, 4 behavioral measures were skewed, so PLS was re-computed after having removed these measures. Fourth, instead of regressing age, sex, education, site and motion from the data, these variables were included with the behavioral data for the PLS analysis. Fifth, instead of GSR in the rs-fMRI preprocessing, we utilized CompCor (53) without GSR. Finally, to ensure our results were not driven by the large number of HC, or by case-control group differences, PLS was re-computed using only controls or only patients.

## Results

### Partial least squares (PLS) reveals three robust latent components linking behavior and brain function

We applied PLS to whole-brain RSFC and 54 behavioral measures of 224 participants across diagnostic categories. **Figure S1** shows the amount of covariance explained by each LC. Four latent components (LCs) survived permutation testing with FDR (q < 0.05) correction. Because LC4 (**Figure S2**) was not robust to control analyses (see below), we focus on LC1 to LC3 for the remainder of this paper. We also note that none of the confounds (age, sex, motion, site and education) examined in **Table 1** were associated with any component (**Table S4**).

### First latent component (LC1) reflects general psychopathology

LC1 accounted for 20% of RSFC-behavior covariance (**Figure S1**) with significant association (r = 0.78, p = 0.007) between RSFC and behavioral composite scores (**Figure 1A**).

**Figure 1.**
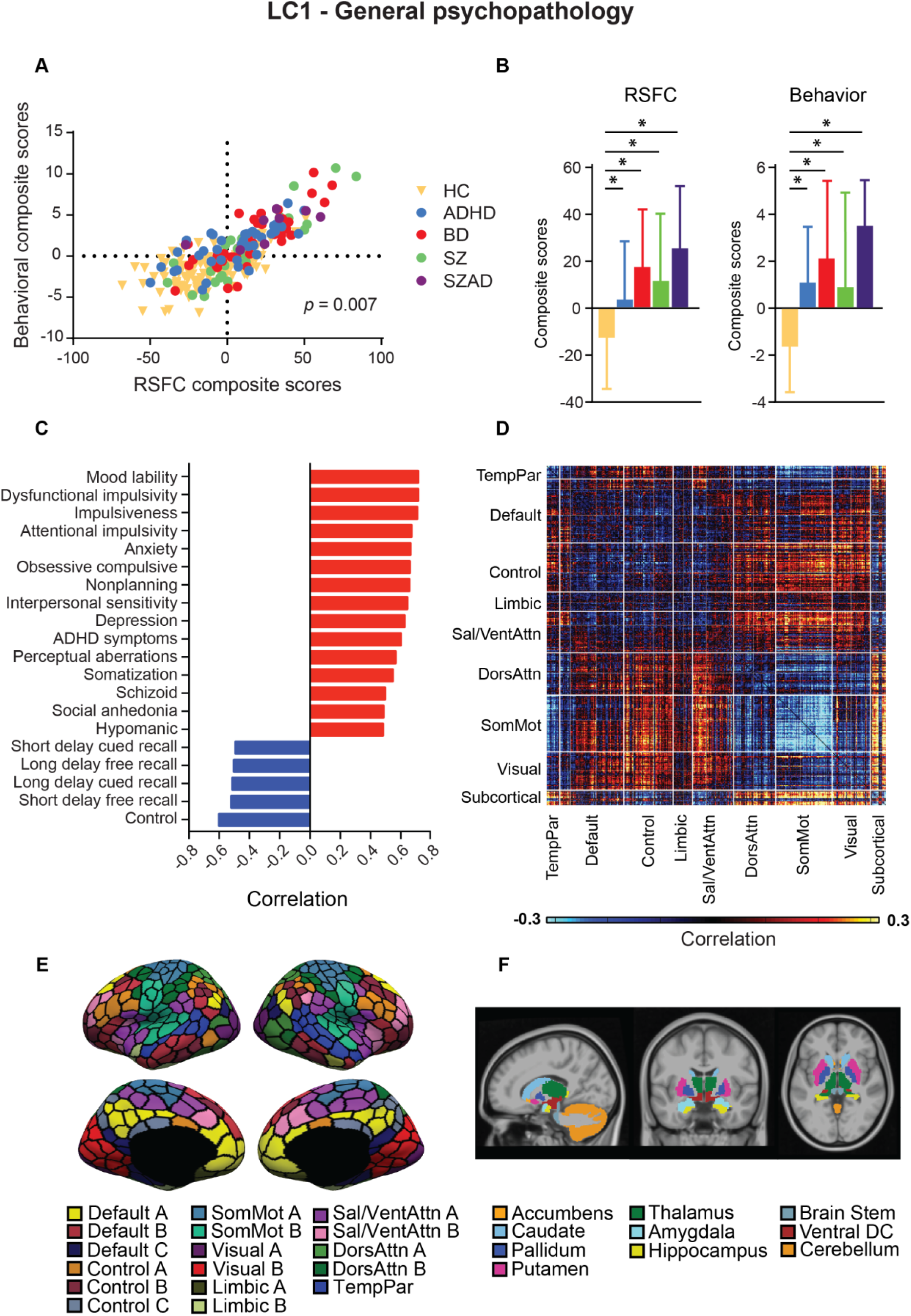
First latent component reflects general psychopathology. (A) Correlation between individual-specific RSFC and behavioral composite scores of participants. Scatterplots for each primary diagnostic group are found in **Figure S11**. (B) Group differences in RSFC and behavioral composite scores. HC had significantly lower RSFC and behavioral composite scores compared to all patient groups. (C) Top 20 strongest correlations between participants’ behavioral measures and their behavioral composite scores. Greater loading on LC1 was associated with higher measures of psychopathology and worse control. (D) Correlations between participants’ RSFC data and their RSFC composite scores. Red (or blue) color indicates that greater RSFC is positively (or negatively) associated with LC1. (E) Schaefer’s 400 cortical regions (47). The parcels are assigned to 17 resting-state networks, which are further grouped into 8 major networks: Temporo-Parietal, Default Mode, Executive Control, Limbic, Salience/Ventral Attention, Dorsal Attention, Somatomotor and Visual (47). (F) 19 subcortical regions (48).

**Figure 1C** shows the top correlations between LC1’s behavioral composite score and the 54 behavioral measures (**Table S5** shows complete table). Greater behavioral composite score was associated with greater psychopathology (e.g., mood lability, dysfunctional impulsivity, anxiety) and worse control (which measures capacity to control one’s behavior).

**Figure 1D** shows the correlations between LC1’s RSFC composite scores and the RSFC data among 400 cortical (**Figure 1E**) and 19 subcortical (**Figure 1F**) ROIs. Greater RSFC composite score was associated with decreased RSFC within the somatosensory-motor (Somatomotor) networks. Sensory-motor (Visual, Somatomotor) and Dorsal Attention networks showed greater RSFC with Control and Salience B networks, and with subcortical regions (thalamus, ventral diencephalon, cerebellum, caudate, putamen, and pallidum).

Consistent with the interpretation that LC1 reflects general psychopathology, both RSFC and behavioral composite scores were lower in controls compared to all patient groups (**Figure 1B**), even after controlling for medication and substance use (**Figure S3**).

Higher RSFC and behavioral composite scores were associated with medication use (**Figure S4**) and greater medication load (**Table S4**). When controlling for diagnosis and substance use, medication targeting the dopaminergic system remained associated with higher behavioral composite scores (**Figure S5**). Individuals using two or more substances also had higher behavioral composite scores than those not using any substance, when controlling for diagnosis and medication use (**Figure S7**).

### Second latent component (LC2) reflects differential cognitive impairment between disorders

LC2 accounted for 12% of RSFC-behavior covariance (**Figure S1**) with significant association (r = 0.83, p = 0.016) between RSFC and behavioral composite scores (**Figure 2A**). Greater behavioral composite score was predominantly associated with worse cognitive performance in language, memory and executive function (**Figure 2C**).

**Figure 2.**
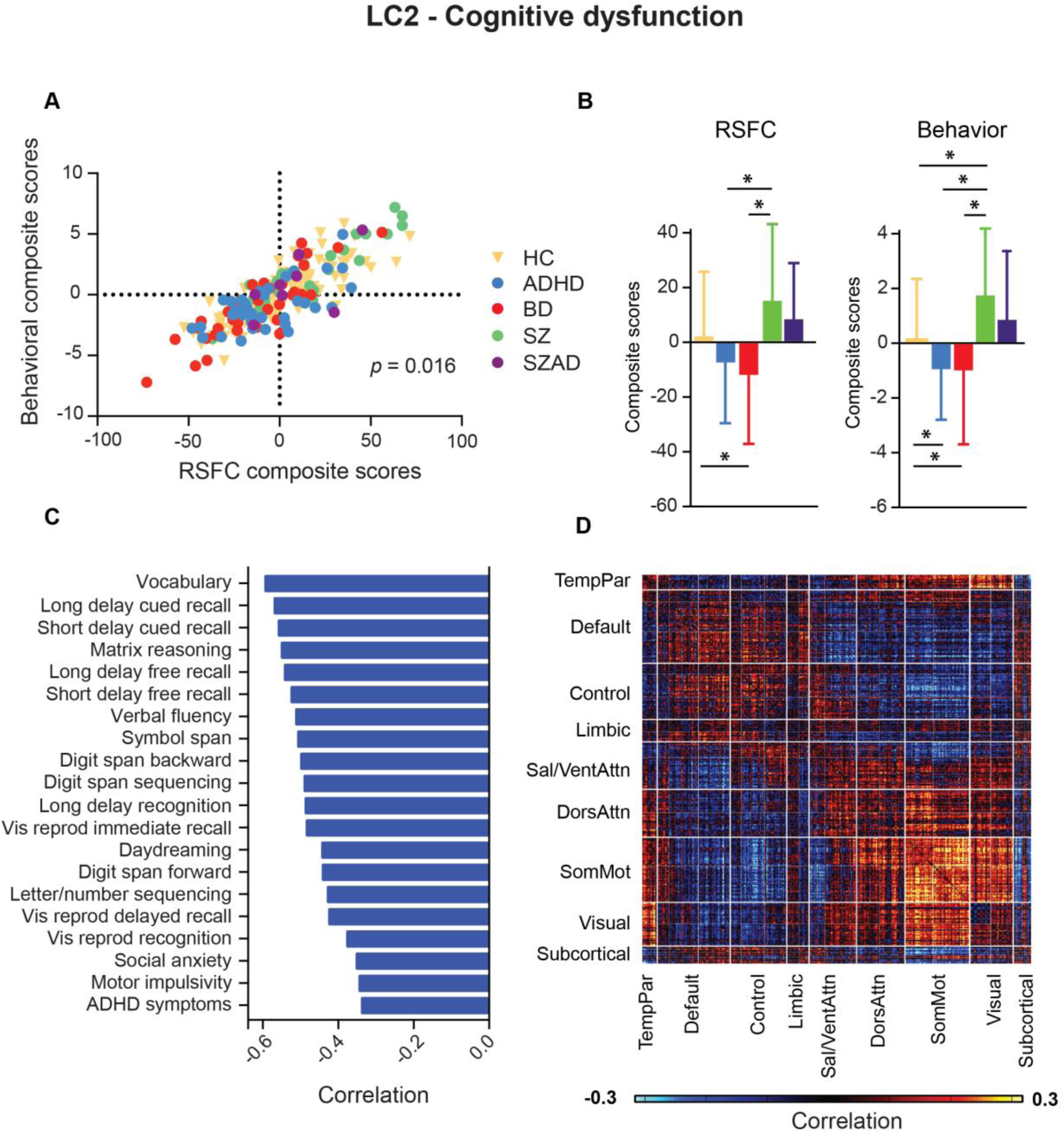
Second latent component reflects cognitive dysfunction. (A) Correlation between individual-specific RSFC and behavioral composite scores of participants. Scatterplots for each primary diagnostic group are found in **Figure S12**. (B) Group differences in RSFC and behavioral composite scores. SZ patients had significantly higher RSFC and behavioral composite scores compared to ADHD and BD. HC also had higher RSFC composite scores than BD, higher behavioral composite scores than ADHD and BD, and lower behavioral composite scores than SZ. (C) Top 20 strongest correlations between participants’ behavioral measures and their behavioral composite scores. Greater loading on LC2 was associated with greater impairment across several cognitive domains, including language, memory and executive functions. (D) Correlations between participants’ RSFC data and their RSFC composite scores. Red (or blue) color indicates that greater RSFC is positively (or negatively) associated with LC2.

Greater RSFC composite score was associated with increased RSFC within the Somatomotor networks (**Figure 2D**). Sensory-motor (Visual, Somatomotor) and attentional (Dorsal Attention, Salience/Ventral Attention) networks showed decreased RSFC with Control and Default networks.

RSFC and behavioral composite scores were higher in SZ/SZAD than in ADHD/BD, although this was only significant with respect to SZ patients (potentially because of the small number of SZAD participants; **Figure 2B**). Differences remained significant after controlling for medication and substance use (**Figure S3**).

Behavioral composite scores were higher in participants taking medication affecting the glutamatergic, norepinephrinergic and dopaminergic systems, as well as other medication, compared to those not taking any medication, when controlling for diagnosis and substance use (**Figure S5**).

Individuals not using any substance had higher RSFC composite scores than those using 2 or more substances (**Figure S6**), but this effect did not survive after controlling for diagnosis and medication use (**Figure S7**).

### Third latent component (LC3) reflects greater impulsivity

LC3 accounted for 8% of RSFC-behavior covariance (**Figure S1**) with significant association (r = 0.73, p = 0.011) between RSFC and behavioral composite scores (**Figure 3A**). **Figure 3C** shows that greater behavioral composite score was associated with greater impulsivity (e.g., functional and motor impulsivity, novelty seeking), as well as lower control, harm avoidance, and social anxiety.

**Figure 3.**
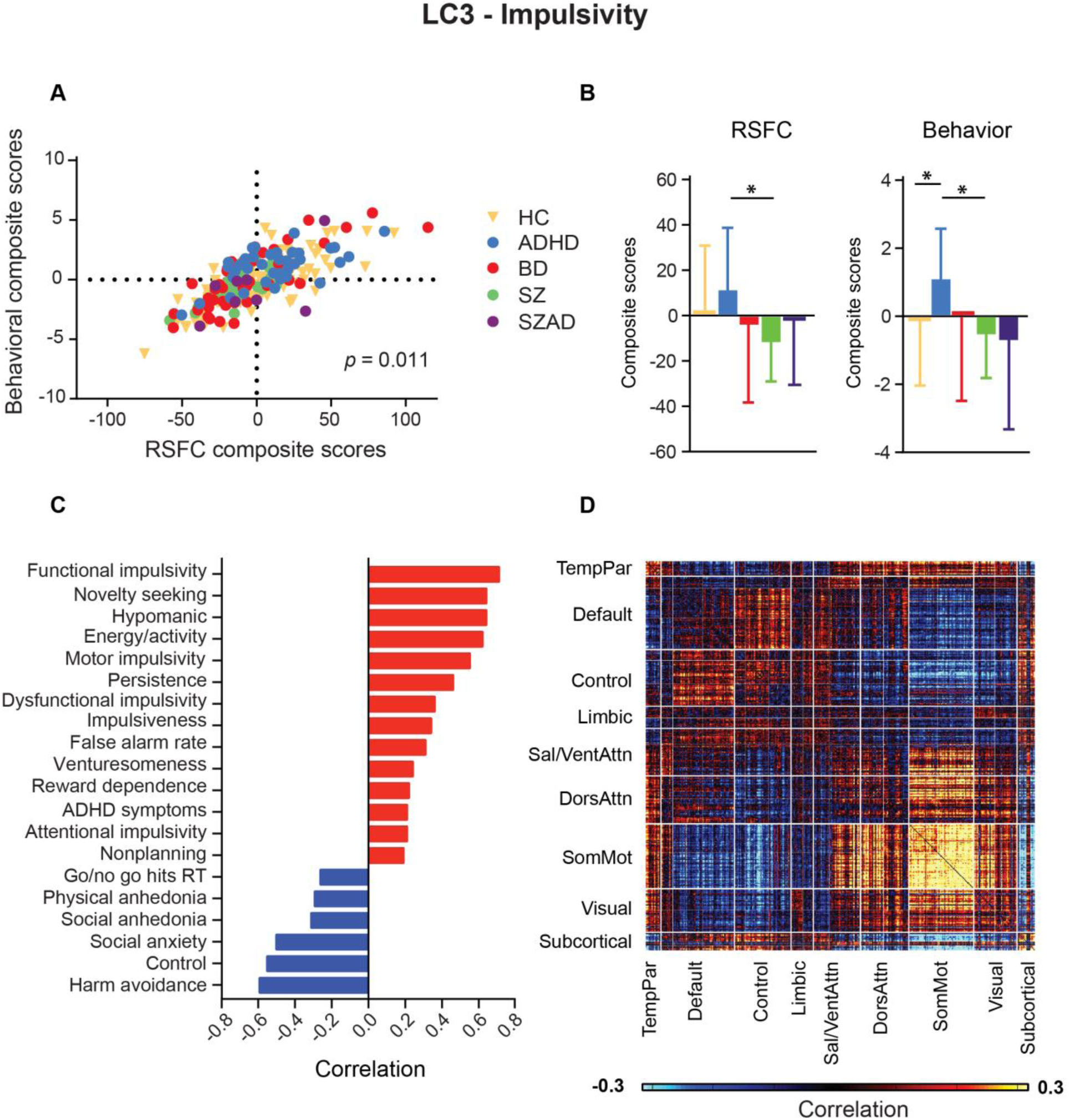
Third latent component reflects impulsivity. (A) Correlation between individual-specific RSFC and behavioral composite scores of participants. Scatterplots for each primary diagnostic group are found in **Figure S13**. (B) Group differences in RSFC and behavioral composite scores. ADHD patients had significantly higher RSFC and behavioral composite scores compared to SZ. ADHD also had significantly higher behavioral composite scores compared to HC. (C) Top 20 strongest correlations between participants’ behavioral measures and their behavioral composite scores. Great loading on LC3 was positively associated with several measures of impulsivity and negatively associated with harm avoidance and control. (D) Correlations between participants’ RSFC data and their RSFC composite scores, showing the connections that contribute most to the LC. Red (or blue) color indicates that greater RSFC is positively (or negatively) associated with LC3.

Greater RSFC composite score was associated with increased RSFC within the Somatomotor networks (**Figure 3D**). Somatomotor networks also showed greater RSFC with Visual B, Dorsal Attention B and Salience A networks, and lower RSFC with Control B network and subcortical regions (caudate, cerebellum and thalamus). Finally, RSFC was also increased between Default (A and B) and Control networks.

ADHD patients had higher RSFC and behavior composite scores than SZ patients, and higher behavioral composite scores than HC (**Figure 3B**). However, only the difference between ADHD and HC was significant after controlling for medication and substance use (**Figure S3**).

Lower RSFC composite score was associated with greater medication load **(Table S4**). Medication affecting the serotonergic, norepinephrinergic and dopaminergic systems were associated with lower RSFC composite scores (**Figure S4**), but these associations did not survive after controlling for diagnosis and substance use (**Figure S5**).

Finally, individuals using one substance had higher RSFC and behavioral composite scores than those not using any substance, and higher behavioral composite scores than those using two or more substances (**Figure S6**), suggesting a nonlinear effect of substance use. The associations with behavioral composite scores remained significant after controlling for diagnosis and medication use (**Figure S7**).

### Somatomotor networks are transdiagnostic hubs

To explore if there was a common set of functional connections across latent components, the absolute correlations between RSFC and RSFC composite scores (**Figures 1D, 2D and 3D**) were thresholded at 0.25 and summed across LCs (**Figure 4**). Somatomotor networks’ connections appeared prominently, including connections within the Somatomotor networks, as well as Somatomotor networks’ connections with Control B, Dorsal Attention B, and subcortical regions (caudate, putamen, thalamus and cerebellum). Results were similar across thresholds (**Figure S8**).

**Figure 4.**
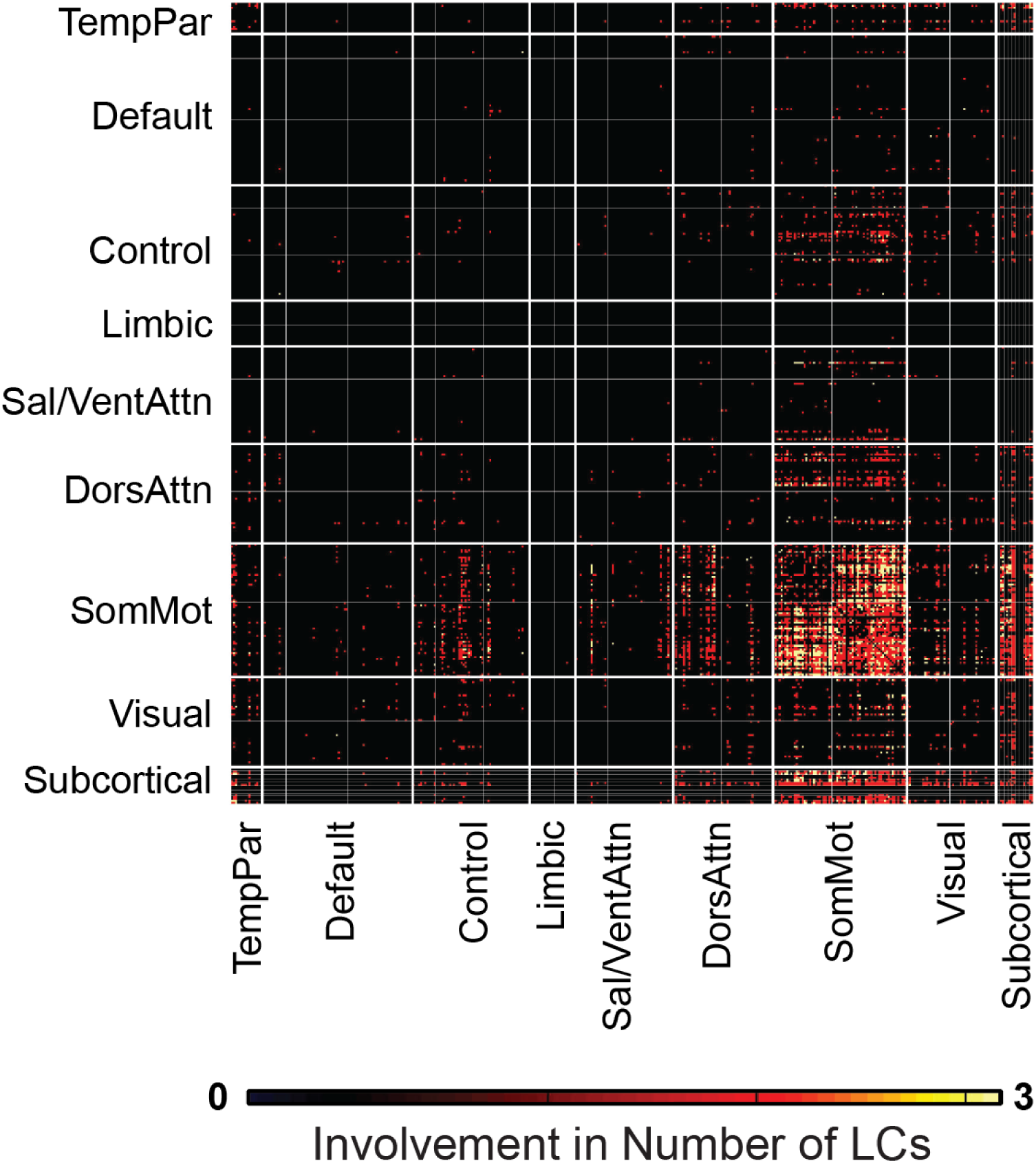
Conjunction map showing connections involved in multiple LCs. Absolute correlations between RSFC and RSFC composite scores (**Figures 1D, 2D and 3D**) were thresholded at 0.25 and summed across LCs. Connections that survived the threshold in only one map were changed to 0 (black); those that survived the threshold in two or three maps had a value of 2 (orange) and 3 (yellow), respectively. Connections involving the Somatomotor networks appeared to be involved in all three LCs. Furthermore, even though we did not focus on LC4 in this study, Somatomotor networks’ connections also featured prominently in LC4 (**Figure S2**).

### Control and reliability analyses

Here, we summarize several analyses that ensured robustness of the LCs. For more details, see **Supplemental Results**. First, PLS components estimated from RSFC successfully generalized to task-FC in the same participants. Second, 5-fold cross-validation was successful: PLS components estimated from 80% of the participants successfully generalized to the remaining 20% of participants. Third, PLS components were robust to certain non-Gaussian and skewed behavioral distributions. Fourth, instead of regressing age, sex, education, site and motion from the data, these variables were included with the behavioral data for the PLS analysis. The results were largely unchanged. Fifth, instead of GSR in the rs-fMRI preprocessing, we utilized CompCor (53) without GSR. The first three LCs were largely unchanged, but not LC4. Hence, we focused on LCs 1-3 in this paper. Finally, to ensure our results were not driven by the large number of HC, or by case-control group differences, PLS was re-computed using only controls or only patients. In both models, we found moderate to high correlations with original saliences. For more details, see **Supplemental Results, Table S6** and **Table S7**.

## Discussion

We identified three latent components representing general psychopathology, cognitive dysfunction and impulsivity, which were associated with distinct whole-brain RSFC patterns across mental health and disease. All three components implicated connectivity of the somatosensory-motor (somatomotor) network with subcortical regions and cortical executive networks. These brain-behavior associations might index intermediate neurobiological processes and potentially serve as transdiagnostic phenotypes, providing a more comprehensive characterization of individuals’ phenotypic variability.

### Somatomotor networks are transdiagnostic hubs

The implication of the somatomotor network across multiple dimensions might seem surprising. However, closer inspection of previous case-control neuroimaging studies suggests that the somatomotor regions are often reported, but not emphasized within prevailing models of psychiatry. For example, altered RSFC within the somatomotor network (54–56), as well as between the somatomotor networks and thalamus have been documented in case-control studies investigating SZ, SZAD and BD patients (54,57–63). Altered thalamo-somatomotor FC has also been linked to SZ symptom severity (54,57,58). One recent study has found RSFC involving somatosensory, motor, basal ganglia, thalamus and visual regions to be associated with psychopathology levels (i.e., *p* factor) in a community-based sample (12). Our results extend previous work by showing that dysconnectivity patterns of these regions are linked to variation in three key domains: general psychopathology, cognitive dysfunction and impulsivity.

In addition to dysconnectivity of the somatomotor network, sensory processing has been found to be disturbed in SZ (64) and BD (65,66). Moreover, the diagnostic criteria for BD and ADHD includes motor features (67,68). Indeed, motor dysfunction has been documented in many psychiatric disorders (69), preceding disease onset and predicting disease progression (70–72). Given that all three LCs in this study featured connectivity between somatomotor and executive networks, these sensory-motor deficits might arise from impaired top-down control over lower-level processes. Another possible mechanism is impaired ability to decode information coming from sensory regions, whereby lower-level sensory deficits may cascade up the system, undermining higher-order cognitive functions (64). Overall, our findings suggest that sensory-motor processes impact symptomatology, cognitive function and personality. Investigating these processes in the future may therefore inform the underlying etiology of various aspects of psychopathology.

### Relations to other transdiagnostic studies

LC1 appeared to reflect the *p* factor widely discussed in transdiagnostic cohorts (10–15). On the other hand, LC2 (cognitive dysfunction) and LC3 (impulsivity) have been featured less frequently and provided further insights into heterogeneity among controls and patients.

LC2 captured dysfunction across multiple cognitive domains. Interestingly, controls had almost no loadings on LC2 (although they generally performed better than patients on all cognitive tests), which argues against LC2 representing general intelligence. Instead, LC2 was largely driven by differences in cognitive performance between SZ/SZAD and ADHD/BD patients. This is consistent with a frontoparietal atrophy pattern being associated with cognitive performance in a transdiagnostic sample, and differentiated between SZ/SZAD and BD patients (73). Previous reports have also suggested similar cognitive deficit patterns among SZ, SZAD and BD patients, although the latter typically exhibited less severe deficits (3,74–77). Overall, this suggests a common etiology underlying general cognitive impairment in these disorders.

Although central to several psychiatric conditions (e.g., ADHD, substance disorder (78)), impulsivity factors have only been reported in one recent transdiagnostic study, but were not associated with any RSFC pattern (12). In our case, the impulsivity measures driving LC3 indexed response inhibition (e.g., false alarm rate) and novelty seeking, and were related to hyperactivity (e.g., energy/activity). This component differentiated ADHD from controls, consistent with hyperactivity/impulsivity being characteristic of ADHD (67).

### Strengths & limitations

One strength of our study is the use of a whole-brain data-driven approach and a broad set of behavioral measures. The components we identified with RSFC generalized well to task fMRI data, and were robust across alternative methodological strategies. Nonetheless, our work has several limitations. First, the sample size of each patient group was small. This was not an issue for the PLS analysis since diagnostic categories were not utilized. However, the limited sample sizes do affect posthoc analyses, e.g., when comparing SZAD loadings with other patient groups. Future research involving larger samples and more diagnostic categories is warranted. Moreover, most scales measuring symptom severity were only administered to patients, which limited the number of clinical measures that could be utilized (c.f. (17)). Finally, our results might be affected by the particular combination of psychiatric disorders and behavioral measures available in this dataset. Future studies will benefit from the increasing availability of broad phenotypic batteries that assess multiple domains of behavior, cognition and genetics (79,80).

## Conclusions

By identifying three components that characterized individuals’ variability in psychopathology, cognitive impairment and impulsivity, our work has allowed to highlight the multifaceted role of somatomotor regions along these dimensions. Our study thus adds further evidence to the benefits of including a broad range of behavioral measures to capture brain-behavior associations across psychiatric boundaries. Identifying such transdiagnostic associations might help uncover common neurobiological mechanisms, and explain high comorbidity rates in psychiatry.

## Acknowledgements

This work was supported by Singapore MOE Tier 2 (MOE2014-T2-2-016), NUS Strategic Research (DPRT/944/09/14), NUS SOM Aspiration Fund (R185000271720), Singapore NMRC (CBRG/0088/2015), NUS YIA and the Singapore National Research Foundation (NRF) Fellowship (Class of 2017). Our research also utilized resources provided by the Center for Functional Neuroimaging Technologies, P41EB015896 and instruments supported by 1S10RR023401, 1S10RR019307, and 1S10RR023043 from the Athinoula A. Martinos Center for Biomedical Imaging at the Massachusetts General Hospital. Our computational work was partially performed on resources of the National Supercomputing Centre, Singapore (https://www.nscc.sg).

All data are publicly available via the OpenfMRI database (https://openfmri.org/dataset/). The accession number of this study is ds000030. We thank the investigators Robert Bilder, Russell Poldrack, Tyrone Cannon, Edythe London, Nelson Freimer, Eliza Congdon, Katherine Karlsgodt and Fred Sabb for sharing their data publicly.

