## Supplementary Material for "Somatosensory-Motor Dysconnectivity Spans Multiple Transdiagnostic Dimensions of Psychopathology"

### *Supplemental Information*

### Supplemental Methods

*Comorbidity*

11 BD patients (out of the original sample, of whom 8 were included in the final sample) had a current comorbid ADHD diagnosis; no other participant had a current comorbid diagnosis that was one of the primary diagnoses (ADHD, BD, SZ, SZAD). Only the primary diagnosis was considered in t-tests assessing the impact of diagnosis on composite scores (**Figure 1B**, **Figure 2B**, **Figure 3B, Figure S2B**), and in ANOVAs assessing group differences in demographic, clinical and behavioral data (**Table 1**, **Table S2**). However, we modelled current comorbid diagnoses (allowing for several concurrent diagnoses in a single individual) in general linear model (GLM) analyses (see below).

*Categorization of medication*

Medication use refers to current use of medication, and was sorted by targeted neurotransmitter system(s), based on the Neuroscience-based Nomenclature (NbN-2 (1,2); http://www.nbn2.com/). Possible targets were the acetylcholinergic, GABAergic, glutamatergic, serotonergic, norepinephrinergic and dopaminergic systems, as well as “other” (which included medication not targeting any neurotransmitter system). Medication targeting the acetylcholinergic system were merged with “other” as both of these categories had too few participants (≤12) on their own. When a medication was not included in this nomenclature, we searched in the psychopharmacological manual Stahl Online (https://stahlonline.cambridge.org/). The list of all medications currently used by participants along with the neurotransmitter system(s) targeted by each medication is presented in **Table S3**.

It is important to note that our assessment of medication was limited. We did not consider dosage, interactions between medications, or the receptor type (agonists/antagonists) targeted by the medication.

As an alternative to neurotransmitter targets, we also utilized a “classical” classification of medication by sorting them into medication classes, which comprised antipsychotics, mood stabilizers, antidepressants, sedatives / hypnotics / anxiolytics (SHA), stimulants, and “other medication” (**Table**

**S3**). Antipsychotics were not further sorted into typical (first generation) and atypical (second generation), as only 4 patients were using typical antipsychotics. The medication classes were tested for association with the RSFC (or behavioral) composite scores using t-tests and also utilized in the GLMs (see below).

*Substance use*

Substance use refers to lifetime substance abuse and/or dependence, and was categorized into abuse and/or dependence on 0/1/2 or more drugs (including nicotine, alcohol, cannabis and other psychotropic substances), resulting in 3 categories. This choice was motivated by having categories with similar number of participants.

*Partial least squares analysis*

Partial least squares (PLS) is a multivariate data-driven statistical technique that aims to extract latent variables (or *latent components* [LCs]) representing maximal brain-behavior associations (3,4). We used an in-house Matlab code from Zoller et al. (5), based on Krishnan et al. (6).

The PLS analysis was computed as follows. The RSFC and behavior data are stored in matrices $X$ (participants $x$ FC values) and $Y$ (participants $x$ behavioral measures). After z-scoring $X$ and $Y$ (across all participants), we computed the covariance matrix $R$:

$$R = Y^T \times X$$

followed by singular value decomposition of $R$:

$$R = U \times S \times V^T$$

which resulted in three low-dimensional matrices: $U$ and $V$ are the singular vectors (called behavioral and RSFC *saliences,* akin to loadings in principal components analysis), while $S$ is a diagonal matrix containing the singular values. Next, we computed $L_X$ and $L_Y$ by projecting $X$ and $Y$ onto their respective saliences $V$ and $U$**:**

$$L_X = X \times V$$

$$L_Y = Y \times U$$

The matrices obtained are the *RSFC* and *behavioral composite scores*, and reflect the participants' individual RSFC and behavioral contribution to each LC (akin to factor scores in principal components analysis). PLS seeks to maximize the covariance between corresponding columns in $L_X$ and $L_Y$. The covariance explained by each LC is estimated by dividing the squared singular value by the sum of all squared singular values (3). The LCs are ordered by the amount of covariance they account for. To assess the statistical significance of the LCs, we permuted the behavioral data (1000 permutations within each primary diagnostic group) and constructed a null distribution of the singular values. To limit the number of multiple comparisons, we only considered the first 5 LCs and applied FDR correction ($q < 0.05$).

In order to infer the contribution of the original variables to the LCs' structure, we computed Pearson's correlations between $X$ and $L_X$, as well as between $Y$ and $L_Y$, yielding *structure coefficients* (7,8). Structure coefficients are commonly computed in factor analysis, multiple regression analysis and canonical correlation analysis (CCA), as they help to define the structure of the synthetic variables (predicted Y scores in factor and regression analyses, canonical function scores in CCA). Indeed, structure coefficients reflect the direct contribution of a predictor to the predictor criterion independently of other predictors, which can be critical when predictors are highly correlated between each other (i.e., in presence of multicollinearity (9)).

Finally, we evaluated the reliability of the RSFC saliences using bootstrap resampling (500 random samples with replacement within each primary diagnostic group), and calculated bootstrap scores for each original RSFC variable by dividing its salience by its bootstrap-estimated standard deviation. Bootstrap scores with absolute values greater than 3 were considered robust at a confidence level of 99% (10). In general, we found the RSFC saliences to be highly similar to the structure coefficients. Correlations between RSFC saliences and structure coefficients for LC1, LC2, LC3 and LC4 were $r = 0.70$, $r = 0.74$, $r = 0.69$ and $r = 0.76$ respectively.

*Replication on task fMRI data*

We tested whether we could replicate the brain-behavior associations identified with rs-fMRI using task fMRI data. We used data from all 7 fMRI tasks (Task Switching, Stop Signal, Paired-Associate Memory (Encoding and Retrieval), Spatial Capacity Working Memory, Balloon Analog Risk and Breath Holding). Task fMRI data were preprocessed using the same pipeline as for rs-fMRI data.

We computed Pearson's correlations between the timeseries of all 419 brain regions in each participant, and then averaged FC of each node of the connectivity matrix across all tasks available for that participant (the median number of tasks available across participants was 6). Averaging FC across all tasks was motivated by an effort to improve signal-to-noise-ratio. Furthermore, some has suggested that task-dependent functional coupling might cluster around a stable state (11,12). By averaging FC across tasks, the resulting FC might be more similar to resting-state FC.

Task FC data were stored in matrix $task\ X$, while behavioral data $task\ Y$ were the same as in the original PLS model (using rs-fMRI data). We z-scored both matrices, and computed $task\ L_X$ and $task\ L_Y$, by projecting $task\ X$ and $task\ Y$ onto their respective saliences of the original PLS model:

$$task\ L_X = task\ X \times rest\ V$$

$$task\ L_Y = task\ Y \times rest\ U$$

Finally, we computed Pearson's correlations between $task\ L_X$ and $task\ L_Y$ for LCs 1-4, and used permutation testing (behavioral data permuted 1000 times within each diagnostic group) to assess whether these correlations were statistically significant.

*Cross-validated PLS*

We carried out a train-test validation of the PLS analysis using *k*-fold cross-validation. First, we assigned 80% of the participants (within each primary diagnostic group) to the training set and the remaining 20% of participants (within each primary diagnostic group) to the test set.

For each fold, we computed PLS on the training dataset $X_{train}$ and $Y_{train}$ , and obtained RSFC and behavioral saliences $V_{train}$ and $U_{train}$. We computed RSFC and behavioral composite scores $L_{X\ train}$ and $L_{Y\ train}$ :

$$L_{X\ train} = X_{train} \times V_{train}$$

$$L_{Y\ train} = Y_{train} \times U_{train}$$

We computed Pearson's correlations between $L_{X\ train}$ and $L_{Y\ train}$ for LCs 1-4 (which were later averaged across the 5 folds).

For each fold, we computed RSFC and behavioral composite scores in the test validation set by projecting the test data $X_{test}$ and $Y_{test}$ onto their respective saliences in the training data:

$$L_{X\ test} = X_{test} \times V_{train}$$

$$L_{Y\ test} = Y_{test} \times U_{train}$$

For each fold, we computed Pearson's correlations between $L_{X\ test}$ and $L_{Y\ test}$ for LCs 1-4, and tested the statistical significance of the correlations using a permutation test (behavioral data permuted 1000 times within each diagnostic group).

*Alternative preprocessing*

Global signal regression (GSR) has been a topic of debate in the neuroimaging community, especially with regards to its impact on RSFC (13,14). Furthermore, patients with psychiatric disorders such as schizophrenia were shown to significantly differ from controls in terms of global fMRI signal (15). Therefore, we considered an alternative preprocessing procedure whereby instead of GSR, we used CompCor (16,17). CompCor derives principal components from nuisance regions-of-interest (ROIs) to model physiological fluctuations that are unlikely to be modulated by neural activity. We used anatomical data to derive white matter and ventricular ROIs, which were reduced to 5 principal components, and then regressed out from the fMRI timeseries. After computing RSFC between all brain regions, we re-computed PLS, and calculated the similarity (using Pearson's correlations) between RSFC (or behavioral) saliences obtained with this model and the RSFC (or behavioral) saliences obtained with the original PLS model.

*Behavior normalization*

Many of the behavioral measures included in the PLS analysis were not Gaussian distributed. To ensure this is not an issue, we applied quantile normalization to all behavioral measures (https://en.wikipedia.org/wiki/Quantile_normalization), and re-computed PLS between RSFC and the normalized behavioral data.

Moreover, 4 behavioral measures (Infrequency, Go/no go hit rate, Long delay recognition and Visual reproduction recognition) were severely left-skewed. We re-computed PLS after having removed these 4 measures.

In both cases, we calculated the similarity between RSFC (or behavioral) saliences obtained with these control analyses and the RSFC (or behavioral) saliences obtained with the original PLS model.

*Confounds*

We regressed out potential confounds such as head motion, age, sex, years of education and acquisition site from both the RSFC and behavior data prior to the PLS analysis. In order to check whether these confounds were associated with the LCs (despite the prior regression), we added them to the behavioral measures (this time without regressing them out of the data). Again, we re-computed PLS, and calculated the similarity between the obtained RSFC (or behavioral) saliences and the RSFC (or behavioral) saliences of the original PLS model. We also checked whether any of these confounds were highly correlated with the behavioral composite scores of the original PLS model.

*Associations between composite scores and diagnosis, medication and substance use*

General linear models (GLMs) were used to test for associations between the RSFC (or behavioral) composite scores and diagnosis (while controlling for medication and substance use). In other words, the RSFC (or behavioral) composite scores were the dependent (response) variables, diagnosis was the independent (predictor) variable, and medication and substance use were added as nuisance covariates. Pairwise comparisons of beta coefficients were then tested using linear hypothesis tests. Associations between participants' diagnosis with RSFC and behavioral composite scores were tested with pairwise comparisons between diagnostic categories (5 variables).

The GLMs were repeated where medication use was the independent (predictor) variable, while diagnosis, substance use and medication load were added as nuisance covariates. Associations between participants' medication use with RSFC and behavioral composite scores were tested with pairwise comparisons between medication categories.

Finally, we considered a separate set of GLMs, where substance use was the independent (predictor) variable, while diagnosis and medication use were added as nuisance covariates. Associations between participants' substance use with RSFC (or behavioral) composite scores were tested with pairwise comparisons between substance use categories.

We note that across all GLMs, both "primary" or "secondary" diagnosis were considered (regardless of whether they were independent or nuisance variables), thus accounting for comorbidity. On the other hand, when used as an independent variable, medication use was coded as 6 variables, one for each targeted neurotransmitter system. When used as a nuisance regressor, we only considered medication load (number of medications used) and intake of *any* medication instead of modeling each neurotransmitter system as a separate regressor in order to limit degrees of freedom. Finally, when used as an independent variable, substance use was categorized into the use of 0/1/2+ substances (3 variables). When used as a nuisance regressor, we considered *any* substance use (as a single regressor) to limit degrees of freedom.

### Supplemental Results

*Control and reliability analyses*

We performed several analyses to ensure robustness of the LCs. First, we showed that the PLS results derived from rs-fMRI generalized to task-fMRI. The behavioral composite scores (which stayed the same in both rs-fMRI and task-fMRI analyses) were significantly correlated with task-FC composite scores with $r = 0.37$ (LC1), 0.40 (LC2), 0.38 (LC3) and 0.42 (LC4). P-values were $< 0.001$ for all LCs (**Table S6**).

Second, 5-fold cross-validation was performed. RSFC and behavioral composite scores for LCs 1-4 were strongly correlated across training folds, with mean correlations across folds between 0.83-0.85. RSFC and behavioral composite scores in the test folds remained significantly correlated (with mean correlations across folds between 0.12-0.18, and p-values ranging from 0.001 to 0.004 (**Table S6**).

Third, we applied quantile normalization to improve the Gaussianity of the behavioral data distributions before PLS. Results were similar to the original PLS, and correlations between the saliences of the new and the original PLS analysis for the first four LCs ranged from 0.93 to 0.99 (**Table S7**). Excluding skewed behavioral measures also yielded similar results (**Table S7**).

Fourth, instead of regressing them from both the RSFC and behavioral data, we added age, sex, years of education, acquisition site and head motion to other behavioral measures for the PLS analysis. Results were similar to the original PLS, with correlations between the saliences of the new and the original PLS analysis ranging from 0.68 to 0.97 for LCs 1-4 (**Table S7**). Furthermore, none of these confounds variables was among the top 20 strongest correlations with behavioral saliences, except for gender (in LC4).

Fifth, instead of using GSR for de-noising the rs-fMRI data, we applied CompCor (16,17). Results were similar to the original PLS. Correlations between the saliences of the new and the original PLS analysis for the first three LCs ranged from 0.75 to 0.96 (see **Table S7**). However, correlations dropped to 0.29-0.56 for LC4; hence we did not describe LC4 further.

We also computed PLS on control participants and on patients only. In the PLS analysis computed using only healthy participants, correlations between the saliences of the new and the original PLS analysis ranged between 0.40-0.80 for the first four LCs. When considering only patients, the first three LCs remained robust (with correlations between 0.68-0.92), but not LC4 (correlations between 0.20-0.22; see **Table S7**).

*Control analyses for medication*

Instead of neurotransmitter targets, the associations between participants' medication use (sorted by medication class) and RSFC (or behavioral) composites scores were tested with t-tests (**Figure S9**). Using any type of medication was associated with higher RSFC and behavioral composite scores in LC1 compared to not using medication. Moreover, using stimulants was associated with lower RSFC composite scores in LC2 compared to antipsychotics as well as with no medication. Behavioral composites scores for LC2 were also higher in individuals using antipsychotics compared to those not using any medication. Finally, RSFC composite scores for LC3 were lower in individuals using antipsychotics and antidepressants compared to those not using medication.

Using GLMs, we also tested for associations between participants' medication use with RSFC (or behavioral) composite scores, while controlling for diagnosis and substance use. However, none of the p-values survived FDR correction (**Figure S10**).

**Table S1**. **Behavioral measures used in the analyses.** This list includes both measures that were used in the PLS analysis and measures that were only considered in posthoc analyses (**Table S5**, marked with *). The number of subjects who had available data for each measure is shown.

| Scale | Subscale/Measure | No. subjects |
|---|---|---|
| Adult Self-Report Scale v.1.1 Screener (18) | ADHD symptoms (total score) | 224 |
| Adult ADHD Clinical Diagnosis Scale* (19) | Inattention<br>Hyperactivity | 114 |
| Hopkins Symptom Checklist (HSCL) (20) | Anxiety<br>Depression<br>Obsessive-compulsiveness<br>Somatization<br>Interpersonal sensitivity | 224 |
| Brief Psychiatric Rating Scale* (21) | Positive symptoms<br>Negative symptoms<br>Mania/disorganization<br>Depression/anxiety | 114 |
| Hamilton Psychiatric Rating Scale for Depression* (HAMD-28) (22) | Total score (items 1-17) | 114 |
| Young Mania Rating Scale-C* (23) | Total score | 114 |
| Scale for the Assessment of Positive Symptoms* (24) | Delusions<br>Hallucinations<br>Bizarre behavior<br>Positive formal thought disorder | 78<br>78<br>78<br>77 |
| Scale for the Assessment of Negative Symptoms* (25) | Alogia<br>Anhedonia<br>Attention<br>Avolition<br>Blunt affect | 78 |
| Chapman Psychosis-Proneness Scales (26) | Perceptual aberrations<br>Social anhedonia<br>Physical anhedonia<br>Infrequency | 224 |
| Scale for Traits that Increase Risk for Bipolar II Disorder | Mood lability<br>Daydreaming<br>Energy/activity<br>Social anxiety | 224 |
| Golden & Meehl's Seven MMPI Items Selected by Taxonomic Method (27) | Schizoid-type personality | 224 |
| Eckblad and Chapman's Hypomanic Personality Scale (28) | Hypomanic personality | 224 |

| Scale | Subscale/Measure | No. subjects |
|---|---|---|
| Temperament and Character Inventory (29) | Reward dependence<br>Persistence<br>Novelty seeking<br>Harm avoidance | 224 |
| Barratt Impulsiveness Scale (BIS-11) (30) | Attentional impulsivity<br>Motor impulsivity<br>Nonplanning | 224 |
| Dickman Impulsivity Scale (31) | Functional impulsivity<br>Dysfunctional impulsivity | 224 |
| Eysenck Impulsivity Inventory (32) | Impulsiveness<br>Venturesomeness<br>Empathy | 224 |
| Multidimensional Personality Questionnaire (33) | Control | 224 |
| Kerby Delay Discounting Task* (34) | Small rewards<br>Medium rewards<br>Large rewards<br>Total | 222 |
| Balloon Analog Risk Task* (35) | Low risk pumps<br>High risk pumps<br>Total pumps | 215 |
| California Verbal Learning Test (CVLT-II) (36) | Short delay free recall<br>Short delay cued recall<br>Long delay free recall<br>Long delay cued recall<br>Long delay recognition | 224 |
| Scene Recognition Task* | Encoding accuracy<br>Encoding RT<br>Recall accuracy<br>Recall RT | 222 |
| Remember-Know Task* | Remember words accuracy | 187 |
| | Remember colors accuracy | 187 |
| | Remember forced recognition 1 feature | 187 |
| | Remember forced recognition 2 features | 187 |
| | Remember mean RT | 184 |
| | Know words accuracy | 187 |
| | Know colors accuracy | 187 |
| | Know forced recognition 1 feature | 187 |
| | Know forced recognition 2 features | 187 |
| | Know mean RT | 179 |
| Wechsler Memory Scale (WMS-IV) (37) | Symbol span<br>Visual reproduction immediate recall<br>Visual reproduction delayed recall<br>Visual reproduction recognition<br>Digit span forward<br>Digit span backward<br>Digit span sequencing | 224 |

| Scale | Subscale/Measure | No. subjects |
|---|---|---|
| Spatial Maintenance and Manipulation Task* (38) | Maintenance mean accuracy<br>Maintenance median RT<br>Manipulation mean accuracy<br>Manipulation median RT | 216 |
| Verbal Maintenance and Manipulation Task* (39) | Maintenance mean accuracy<br>Maintenance median RT<br>Manipulation mean accuracy<br>Manipulation median RT | 213 |
| Spatial Capacity Task* (40) | Load 1 accuracy<br>Load 1 mean RT<br>Load 3 accuracy<br>Load 3 mean RT<br>Load 5 accuracy<br>Load 5 mean RT<br>Load 7 accuracy<br>Load 7 mean RT<br>Maximum capacity | 222 |
| Verbal Capacity Task* (41) | Load 3 accuracy<br>Load 3 mean RT<br>Load 5 accuracy<br>Load 5 mean RT<br>Load 7 accuracy<br>Load 7 mean RT<br>Load 9 accuracy<br>Load 9 mean RT<br>Maximum capacity | 223 |
| Wechsler Adult Intelligence Scale (WAIS-IV) (42) | Matrix reasoning<br>Letter/number sequencing<br>Vocabulary | 224 |
| Stroop Color Word Task* (43) | Interference accuracy<br>Interference RT | 223 |
| Color Trail Test (30) | Interference index | 224 |
| Stop Signal Task* | Quantile RT | 223 |
| Task Switching Task (44,45) | Accuracy<br>Interference<br>Switching cost<br>Residual switching cost | 224 |
| Attention Network Task (46) | Interference RT | 224 |
| Continuous Performance Go/No Go Task (47) | Hit rate<br>Hits median RT<br>False alarm rate | 224 |
| Delis-Kaplan Executive Function System (48) | English verbal fluency<br>Spanish verbal fluency* | 224<br>78 |

*Abbreviations*: ADHD, attention deficit/hyperactivity disorder; MMPI, Minnesota Multiphasic Personality Inventory; RT, reaction time.

**Table S2. Group scores on all behavioral measures included in the PLS analysis**. Mean (standard deviation) values are shown for each primary diagnostic group. Groups were compared with ANOVAs.

| Scale / Subscale | | HC | ADHD | BD | SZ | SZAD | *F* | *p* value |
|---|---|---|---|---|---|---|---|---|
| Hopkins Symptom Checklist | Depression | 0.37 (0.36) | 0.67 (0.47) | 0.91 (0.64) | 0.69 (0.56) | 1.19 (0.63) | 14.16 | **2.7E-10** |
| | Obsessive compulsiveness | 0.50 (0.43) | 1.21 (0.70) | 1.13 (0.77) | 0.96 (0.56) | 1.20 (0.76) | 16.84 | **4.7E-12** |
| | Anxiety | 0.20 (0.29) | 0.46 (0.41) | 0.73 (0.65) | 0.70 (0.67) | 0.94 (0.54) | 16.12 | **1.4E-11** |
| | Somatization | 0.21 (0.23) | 0.36 (0.28) | 0.64 (0.66) | 0.54 (0.44) | 0.93 (0.48) | 14.87 | **9.1E-11** |
| | Interpersonal sensitivity | 0.40 (0.36) | 0.77 (0.52) | 1.05 (0.75) | 0.86 (0.65) | 1.02 (0.66) | 14.56 | **1.5E-10** |
| Adult Self Report Scale | ADHD symptoms | 8.67 (2.78) | 15.38 (3.83) | 13.05 (4.96) | 8.90 (4.26) | 11.38 (4.21) | 29.24 | **1.74E-19** |
| Barratt Impulsiveness Scale | Attentional impulsivity | 14.31 (3.35) | 21.54 (4.19) | 19.45 (5.39) | 16.28 (4.44) | 18.63 (4.47) | 27.20 | **2.5E-18** |
| | Motor impulsivity | 21.60 (3.74) | 26.54 (3.96) | 24.60 (5.32) | 21.79 (3.92) | 25.38 (5.13) | 12.46 | **3.8E-09** |
| | Nonplanning | 22.74 (4.37) | 28.22 (4.27) | 28.80 (5.90) | 25.24 (5.80) | 28.00 (2.45) | 17.07 | **3.3E-12** |
| Dickman Impulsivity Scale | Functional impulsivity | 6.49 (2.66) | 6.73 (2.85) | 5.63 (3.35) | 5.55 (2.54) | 5.38 (3.66) | 1.56 | 1.9E-01 |
| | Dysfunctional impulsivity | 1.79 (2.38) | 4.73 (2.88) | 5.93 (4.46) | 3.45 (3.26) | 5.13 (2.80) | 16.83 | **4.8E-12** |
| Multidimensional Personality Questionnaire | Control | 18.38 (4.83) | 11.24 (4.87) | 11.03 (7.40) | 16.83 (4.23) | 14.63 (5.73) | 21.35 | **7.0E-15** |
| Eysenck Impulsivity Inventory | Impulsiveness | 6.02 (2.91) | 9.03 (3.28) | 9.35 (4.43) | 8.93 (3.38) | 10.12 (4.02) | 12.56 | **3.2E-09** |
| | Venturesome-ness | 8.63 (2.44) | 9.35 (1.84) | 7.95 (2.46) | 8.07 (2.78) | 8.38 (2.07) | 2.00 | 9.6E-02 |
| | Empathy | 10.61 (3.04) | 11.35 (2.97) | 11.03 (3.43) | 10.21 (3.08) | 11.75 (2.25) | 0.91 | 4.6E-01 |
| Scale for Traits that Increase Risk for Bipolar II Disorder | Mood lability | 2.06 (1.71) | 3.84 (2.41) | 5.45 (2.94) | 3.07 (2.40) | 6.38 (2.00) | 22.84 | **8.7E-16** |
| | Energy/activity | 3.04 (2.11) | 3.59 (1.99) | 3.95 (2.51) | 4.45 (2.05) | 3.12 (1.96) | 3.18 | 1.4E-02 |
| | Daydreaming | 3.02 (1.79) | 3.97 (1.28) | 3.65 (1.81) | 2.76 (2.08) | 4.38 (1.51) | 3.93 | **4.2E-03** |
| | Social anxiety | 3.01 (1.79) | 3.22 (1.72) | 3.70 (2.00) | 3.55 (1.74) | 3.62 (1.85) | 1.38 | 2.4E-01 |
| Golden & Meehl's Seven MMPI | Schizoid personality | 2.45 (1.26) | 3.05 (1.31) | 4.03 (1.67) | 3.17 (1.81) | 4.50 (0.93) | 11.74 | **1.2E-08** |
| Eckblad and Chapman's Hypomanic Personality Scale | Hypomanic personality | 16.27 (7.69) | 24.46 (7.35) | 23.93 (11.22) | 19.86 (8.68) | 19.88 (8.79) | 9.76 | **2.8E-07** |

| Scale / Subscale | | HC | ADHD | BD | SZ | SZAD | *F* | *p* value |
|---|---|---|---|---|---|---|---|---|
| Chapman Psychosis-Proneness Scales | Infrequency | 0.65 (1.07) | 0.78 (1.08) | 0.85 (1.17) | 1.48 (1.43) | 2.38 (2.07) | 6.13 | **1.1E-04** |
| | Perceptual aberrations | 2.05 (2.54) | 4.00 (3.75) | 4.97 (4.56) | 8.48 (8.46) | 13.25 (4.98) | 22.46 | **1.5E-15** |
| | Social anhedonia | 9.73 (6.79) | 13.95 (8.72) | 16.07 (7.28) | 14.24 (6.26) | 17.38 (6.95) | 8.32 | **2.9E-06** |
| | Physical anhedonia | 11.10 (6.48) | 13.14 (7.79) | 15.97 (9.41) | 15.83 (6.63) | 14.62 (7.46) | 4.65 | **1.3E-03** |
| Temperament and Character Inventory | Persistence | 24.23 (6.92) | 21.43 (7.57) | 18.60 (9.63) | 22.55 (6.69) | 18.62 (6.09) | 4.83 | **9.5E-04** |
| | Harm avoidance | 11.49 (6.26) | 12.54 (6.25) | 18.55 (9.11) | 13.41 (6.31) | 21.25 (8.00) | 10.34 | **1.1E-07** |
| | Reward dependence | 16.12 (4.18) | 14.59 (4.80) | 13.85 (5.22) | 14.07 (4.23) | 14.25 (3.06) | 2.79 | 2.7E-02 |
| | Novelty seeking | 18.91 (5.88) | 24.78 (4.82) | 23.23 (7.77) | 18.17 (4.79) | 19.12 (6.17) | 9.92 | **2.2E-07** |
| Task Switching | Total accuracy | 0.97 (0.03) | 0.96 (0.03) | 0.96 (0.03) | 0.93 (0.07) | 0.93 (0.03) | 7.46 | **1.2E-05** |
| | Interference | 43.18 (61.31) | 55.32 (88.62) | 67.08 (74.23) | 98.90 (206.79) | 50.81 (118.01) | 1.90 | 1.1E-01 |
| | Switch cost | 263.52 (143.74) | 273.23 (125.40) | 243.66 (134.29) | 298.69 (252.47) | 329.38 (148.25) | 0.85 | 5.0E-01 |
| | Residual switch cost | 57.81 (100.58) | 84.73 (121.94) | 50.59 (104.10) | 83.95 (212.75) | 112.38 (62.61) | 0.92 | 4.5E-01 |
| Continuous Performance Go/NoGo Task | Hit rate | 0.99 (0.01) | 0.99 (0.02) | 0.99 (0.03) | 0.98 (0.04) | 0.97 (0.04) | 3.90 | **4.4E-03** |
| | False alarm rate | 0.35 (0.18) | 0.40 (0.22) | 0.36 (0.21) | 0.36 (0.16) | 0.46 (0.24) | 0.88 | 4.8E-01 |
| | Hits RT | 353.92 (44.05) | 358.39 (51.55) | 389.65 (62.18) | 396.59 (47.86) | 375.75 (58.19) | 6.83 | **3.4E-05** |
| California Verbal Learning Test | Short delay free recall | 12.95 (2.30) | 11.76 (2.77) | 10.55 (3.39) | 8.66 (3.68) | 8.25 (2.87) | 18.55 | **3.8E-13** |
| | Short delay cued recall | 13.41 (2.04) | 12.35 (2.30) | 11.65 (3.07) | 9.62 (2.97) | 10.12 (2.10) | 16.93 | **4.1E-12** |
| | Long delay free recall | 13.26 (2.34) | 12.30 (2.39) | 11.05 (3.27) | 9.45 (3.08) | 9.25 (3.06) | 16.37 | **9.6E-12** |
| | Long delay cued recall | 13.69 (2.02) | 12.76 (2.19) | 11.88 (3.41) | 9.90 (3.21) | 9.88 (2.53) | 16.54 | **7.4E-12** |
| | Long delay recognition | 3.37 (0.82) | 3.32 (0.63) | 3.25 (0.87) | 2.58 (0.96) | 2.41 (1.17) | 7.19 | **1.9E-05** |
| Wechsler Memory Scale | Visual reproduction immediate recall | 38.25 (4.60) | 37.54 (5.69) | 35.77 (4.84) | 31.86 (8.42) | 34.25 (8.19) | 8.42 | **2.5E-06** |
| | Visual reproduction delayed recall | 32.79 (8.50) | 29.97 (9.37) | 26.85 (10.26) | 23.90 (10.92) | 23.62 (11.34) | 7.48 | **1.2E-05** |
| | Visual reproduction recognition | 6.49 (0.78) | 6.51 (0.87) | 6.20 (1.07) | 5.52 (1.68) | 6.00 (0.93) | 6.05 | **1.2E-04** |

| Scale / Subscale | | HC | ADHD | BD | SZ | SZAD | *F* | *p* value |
|---|---|---|---|---|---|---|---|---|
| Wechsler Memory Scale | Symbol span | 25.74 (6.34) | 23.14 (6.73) | 21.77 (6.48) | 16.93 (5.50) | 19.75 (9.51) | 12.14 | **6.3E-09** |
| | Digit span forward | 10.92 (2.49) | 11.16 (2.06) | 10.72 (2.32) | 9.17 (2.14) | 8.25 (1.58) | 5.87 | **1.7E-04** |
| | Digit span backward | 9.49 (2.53) | 9.03 (2.23) | 8.93 (2.35) | 7.55 (2.03) | 6.88 (2.47) | 5.41 | **3.6E-04** |
| | Digit span sequencing | 9.74 (2.30) | 9.16 (1.86) | 8.72 (2.63) | 7.59 (1.82) | 7.38 (1.92) | 7.05 | **2.3E-05** |
| Wechsler Adult Intelligence Scale | Letter/number sequencing | 21.05 (2.87) | 20.05 (2.68) | 19.65 (2.57) | 18.55 (2.78) | 16.12 (4.73) | 9.32 | **5.7E-07** |
| | Vocabulary | 43.75 (8.25) | 43.46 (9.65) | 42.92 (10.37) | 30.48 (9.43) | 36.38 (8.40) | 13.75 | **5.1E-10** |
| | Matrix reasoning | 20.65 (4.07) | 20.70 (3.96) | 19.62 (4.42) | 14.79 (4.89) | 17.62 (4.72) | 11.97 | **8.1E-09** |
| Color Trails Test | Interference | 1.12 (0.54) | 1.06 (0.64) | 1.13 (0.57) | 1.19 (0.80) | 0.92 (0.47) | 0.41 | 8.0E-01 |
| Attention Network Task | Interference | 79.39 (32.76) | 81.01 (36.34) | 73.31 (38.25) | 77.18 (41.71) | 105.15 (62.97) | 1.29 | 2.8E-01 |
| English Verbal fluency | Letter fluency | 41.55 (12.74) | 40.84 (10.31) | 40.05 (13.88) | 30.03 (8.10) | 28.50 (8.83) | 7.12 | **2.1E-05** |

All p-values that survived FDR correction ($q < 0.05$) are indicated in bold.
*Abbreviations*: HC, healthy controls; ADHD, attention deficit/hyperactivity disorder; BD, bipolar disorder; SZ, schizophrenia; SZAD, schizoaffective disorder; MMPI, Minnesota Multiphasic Personality Inventory; RT, reaction time.

**Table S3. Categorization of medication by targeted neurotransmitter system(s) and medication class**

| Medication / Molecule | Targeted neurotransmitter system(s) | Class |
|---|---|---|
| Abilify / Aripiprazole | Serotonin, Dopamine | Antipsychotic |
| Adapin, Sinequan / Doxepin | Serotonin, Norepinephrine | Antidepressant |
| Adderall / Dextroamphetamine | Norepinephrine, Dopamine | Stimulant |
| Ambien / Zolpidem | GABA | SHA |
| Anafranil / Clomipramine | Serotonin, Norepinephrine | Antidepressant |
| Artane / Trihexyphenidyl | Acetylcholine | Other |
| Atapryl, Carbex, Eldepryl / Selegiline | Serotonin, Norepinephrine, Dopamine | Other |
| Atarax, Vistaril / Hydroxyzine | Other | SHA |
| Ativan / Lorazepam | GABA | SHA |
| Benadryl / Diphenhydramine | Other | Other |
| Buspar / Buspirone | Serotonin | SHA |
| Celexa / Citalopram | Serotonin | Antidepressant |
| Clozaril / Clozapine | Serotonin, Norepinephrine, Dopamine | Antipsychotic |
| Cogentin / Benzatropine | Acetylcholine, Dopamine | Other |
| Concerta / Methylphenidate XR | Norepinephrine, Dopamine | Stimulant |
| Darvocet / Propoxyphene | Other | Other |
| Depakote / Divalproex | GABA | Mood stabilizer |
| Desyrel / Trazodone | Serotonin | Antidepressant |
| Dexedrine / Dextroamphetamine | Norepinephrine, Dopamine | Stimulant |
| Effexor / Venlafaxine | Serotonin, Norepinephrine | Antidepressant |
| Elavil, Endep / Amitriptyline | Serotonin, Norepinephrine | Antidepressant |
| Eskalith, Lithonate / Lithium Carbonate | Glutamate | Mood stabilizer |
| Fanapt / Iloperidone | Serotonin, Dopamine | Antipsychotic |
| Geodon / Ziprasidone | Serotonin, Dopamine | Antipsychotic |
| Haldol / Haloperidol | Dopamine | Antipsychotic |
| Inderal / Propranolol | Other | Other |
| Invega / Paliperidone | Serotonin, Norepinephrine, Dopamine | Antipsychotic |
| Klonopin / Clonazepam | GABA | SHA |
| Lamictal / Lamotrigine | Glutamate | Mood stabilizer |
| Levothroid / Levothyroxine sodium | Other | Other |
| Lexapro / Escitalopram oxalate | Serotonin | Antidepressant |
| Loxitane / Loxapine | Serotonin, Dopamine | Antipsychotic |
| Metadate / Methylphenidate XR | Norepinephrine, Dopamine | Stimulant |
| Navane / Thiothixene | Dopamine | Antipsychotic |
| Neurontin / Gabapentin | Glutamate | Mood stabilizer |
| Paxil / Paroxetine | Serotonin | Antidepressant |
| Phentermine Hydrochloride | Norepinephrine, Dopamine | Stimulant |
| Prolixin / Fluphenazine | Dopamine | Antipsychotic |
| Provigil / Modafinil | Dopamine | Stimulant |
| Prozac / Fluoxetine | Serotonin | Antidepressant |
| Remeron / Mirtazapine | Serotonin, Norepinephrine | Antidepressant |
| Restoril / Temazepam | GABA | SHA |
| Risperdal / Risperidone | Serotonin, Norepinephrine, Dopamine | Antipsychotic |
| Ritalin / Methylphenidate | Norepinephrine, Dopamine | Stimulant |
| Rozerem / Ramelteon | Other | SHA |
| Saphris / Asenapine | Serotonin, Norepinephrine, Dopamine | Antipsychotic |
| Seroquel / Quetiapine | Serotonin, Dopamine | Antipsychotic |
| Sleep Eze / Diphenhydramine | Other | SHA |
| Synthroid / Levothyroxine sodium | Other | Other |
| Tegretol / Carbamazepine | Glutamate | Mood stabilizer |

| **Medication / Molecule** | **Targeted neurotransmitter system(s)** | **Class** |
|---|---|---|
| Tenormin / Atenolol | Other | Other |
| Topamax / Topiramate | GABA, Glutamate | Mood stabilizer |
| Trileptal / Oxcarbazepine | Glutamate | Mood stabilizer |
| Vicodin / Hydrocodone+ acetaminophen | Other | Other |
| Vyvanse / Lixdexamfetamine | Norepinephrine, Dopamine | Stimulant |
| Wellbutrin / Bupropion | Norepinephrine, Dopamine | Antidepressant |
| Xanax / Alprazolam | GABA | SHA |
| Xyrem / Sodium oxybate | GABA | Other |
| Zoloft / Sertraline | Serotonin | Antidepressant |
| Zyprexa / Olanzapine | Serotonin, Dopamine | Antipsychotic |

Possible neurotransmitter targets were the acetylcholinergic, GABAergic, glutamatergic, serotonergic, norepinephrinergic, and dopaminergic systems, as well as “other” (which included medication not targeting any neurotransmitter system). Medication targeting the acetylcholinergic system were merged with “other medication” as both of these categories had too few participants (≤12) on their own. Medication classes comprised antipsychotics, mood stabilizers, antidepressants, sedatives/hypnotics/anxiolytics (SHA), stimulants, and “other”. Electroconvulsive therapy was categorized as “other” for both targeted neurotransmitter system and medication class.

**Table S4. Associations between RSFC (or behavioral) composite scores and confounds**. Either Pearson's correlations (for continuous measures) or t-tests (for categorical measures) were computed across all participants.

| | LC1 | | | | LC2 | | | | LC3 | | | |
|---|---|---|---|---|---|---|---|---|---|---|---|---|
| | RSFC composite scores | | Behavioral composite scores | | RSFC composite scores | | Behavioral composite scores | | RSFC composite scores | | Behavioral composite scores | |
| | *r / t* | *p* | *r / t* | *p* | *r / t* | *p* | *r / t* | *p* | *r / t* | *p* | *r / t* | *p* |
| Age | 0.00 | 1.000 | 0.00 | 1.000 | 0.00 | 1.000 | 0.00 | 1.000 | 0.00 | 1.000 | 0.00 | 1.000 |
| Gender | 0.00 | 1.000 | 0.00 | 1.000 | 0.00 | 1.000 | 0.00 | 1.000 | 0.00 | 1.000 | 0.00 | 1.000 |
| Education | 0.00 | 1.000 | 0.00 | 1.000 | 0.00 | 1.000 | 0.00 | 1.000 | 0.00 | 1.000 | 0.00 | 1.000 |
| Site | 0.00 | 1.000 | 0.00 | 1.000 | 0.00 | 1.000 | 0.00 | 1.000 | 0.00 | 1.000 | 0.00 | 1.000 |
| Head motion | 0.00 | 1.000 | 0.00 | 1.000 | 0.00 | 1.000 | 0.00 | 1.000 | 0.00 | 1.000 | 0.00 | 1.000 |
| # meds | **0.48** | **1.7E-14** | **0.47** | **1.1E-13** | -0.02 | 0.799 | 0.08 | 0.243 | **-0.21** | **0.002** | -0.13 | 0.047 |

Associations that survived FDR correction (q < 0.05) are indicated in bold.

**Table S5. Correlations between participants' behavioral measures and their behavioral composite scores**. These correlations (sorted in decreasing order) show the contribution of each behavioral measure to latent components (LCs) 1-4. This list includes both measures that were available for all participants (and thus included in the PLS analysis) and measures with missing data (marked with *, not included in the PLS analysis).

| LC1 | | LC2 | | LC3 | | LC4 | |
|---|---|---|---|---|---|---|---|
| **Behavior** | **Corr** | **Behavior** | **Corr** | **Behavior** | **Corr** | **Behavior** | **Corr** |
| Mood lability | 0.72 | Infrequency | 0.29 | Functional impulsivity | 0.71 | Somatization | 0.46 |
| Dysfunctional impulsivity | 0.71 | Alogia* | 0.28 | Novelty seeking | 0.64 | Visual reproduction delayed recall | 0.41 |
| Impulsiveness | 0.71 | Control | 0.28 | Hypomanic personality | 0.64 | Depression/ anxiety* | 0.40 |
| Attentional impulsivity | 0.67 | Blunt affect* | 0.26 | Energy/ activity | 0.62 | Anxiety | 0.40 |
| Anxiety | 0.66 | Attention* | 0.21 | Motor impulsivity | 0.55 | Depression (HSCL) | 0.39 |
| Obsessive compulsive | 0.66 | Hallucinations* | 0.19 | Persistence | 0.46 | Perceptual aberrations | 0.36 |
| Nonplanning | 0.66 | Go/no go false alarm rate | 0.18 | Dysfunctional impulsivity | 0.36 | Energy/ activity | 0.36 |
| Interpersonal sensitivity | 0.64 | Persistence | 0.17 | Hyperactivity* | 0.34 | Interpersonal sensitivity | 0.35 |
| Depression (HSCL) | 0.63 | Know RT* | 0.16 | Impulsiveness | 0.34 | Persistence | 0.35 |
| ADHD symptoms | 0.60 | Stop signal RT* | 0.16 | Go/no go false alarm rate | 0.31 | Infrequency | 0.35 |
| Perceptual aberrations | 0.57 | Attention network task interference RT | 0.14 | Inattention* | 0.26 | Anhedonia* | 0.32 |
| Somatization | 0.55 | Negative symptoms* | 0.14 | Venturesomeness | 0.24 | Depression (HAMD)* | 0.31 |
| Schizoid personality | 0.50 | Spatial WM load 1 RT* | 0.13 | Reward dependence | 0.22 | Obsessive compulsive | 0.31 |
| Social anhedonia | 0.49 | Spatial WM load 7 RT* | 0.13 | ADHD symptoms | 0.21 | Visual reproduction immediate recall | 0.30 |
| Hypomanic personality | 0.48 | Spatial WM load 3 RT* | 0.12 | Attentional impulsivity | 0.21 | Physical anhedonia | 0.28 |
| Harm avoidance | 0.48 | Spatial maintenance RT* | 0.10 | Nonplanning | 0.19 | Spanish verbal fluency* | 0.27 |
| Motor impulsivity | 0.46 | Delusions* | 0.09 | Mania/ disorganization* | 0.17 | Positive symptoms* | 0.26 |
| Social anxiety | 0.42 | Positive symptoms* | 0.09 | English verbal fluency | 0.15 | Hypomanic personality | 0.26 |
| Physical anhedonia | 0.41 | Verbal maintenance RT* | 0.09 | Color trail interference | 0.14 | Social anhedonia | 0.23 |
| Daydreaming | 0.36 | Color trail interference | 0.08 | Bizarre behavior* | 0.12 | Negative symptoms* | 0.22 |
| Depression (HAMD)* | 0.34 | Spatial WM load 5 RT* | 0.08 | Empathy | 0.11 | Mania* | 0.22 |
| Depression/ anxiety* | 0.33 | Task switching interference | 0.07 | Verbal maintenance accuracy* | 0.11 | Long delay cued recall | 0.21 |
| Novelty seeking | 0.32 | Scene recognition encoding RT* | 0.07 | Stroop accuracy* | 0.11 | Short delay cued recall | 0.21 |
| Infrequency | 0.32 | Verbal WM load 5 RT* | 0.06 | Visual reproduction recognition | 0.11 | Bizarre behavior* | 0.21 |
| Empathy | 0.31 | Delay discounting small rewards* | 0.06 | Digit span sequencing | 0.10 | Delusions* | 0.20 |
| Delusions* | 0.28 | Delay discounting total* | 0.06 | Verbal WM load 3 accuracy* | 0.10 | Hallucinations* | 0.20 |
| Energy/ activity | 0.26 | Task switching switch cost | 0.06 | Know forced recognition 2 features* | 0.10 | Long delay free recall | 0.20 |

| LC1 | | LC2 | | LC3 | | LC4 | |
|---|---|---|---|---|---|---|---|
| **Behavior** | **Corr** | **Behavior** | **Corr** | **Behavior** | **Corr** | **Behavior** | **Corr** |
| Positive symptoms* | 0.25 | Delay discounting medium rewards* | 0.06 | Digit span forward | 0.10 | Mood lability | 0.17 |
| Avolition* | 0.22 | Delay discounting large rewards* | 0.05 | Positive formal thought disorder* | 0.09 | Visual reproduction recognition | 0.15 |
| Anhedonia* | 0.21 | Verbal WM load 3 RT* | 0.05 | Verbal WM load 7 accuracy* | 0.09 | Mania/ disorganization* | 0.14 |
| Inattention* | 0.21 | Perceptual aberrations | 0.04 | Matrix reasoning | 0.09 | Positive formal thought disorder* | 0.14 |
| Hallucinations* | 0.19 | Go/no go hit rate | 0.03 | Letter/number sequencing | 0.09 | Spatial maintenance accuracy* | 0.14 |
| Bizarre behavior* | 0.19 | Avolition* | 0.03 | Delay discounting small rewards* | 0.08 | Short delay free recall | 0.13 |
| Alogia* | 0.19 | Verbal manipulation RT* | 0.03 | Know accuracy colors* | 0.08 | Remember RT* | 0.12 |
| Negative symptoms* | 0.18 | Spatial manipulation RT* | 0.02 | Verbal manipulation RT* | 0.07 | Blunt affect* | 0.12 |
| Blunt affect* | 0.17 | Scene recognition recall RT* | 0.02 | Spanish verbal fluency* | 0.07 | Social anxiety | 0.12 |
| Mania* | 0.16 | Physical anhedonia | 0.02 | Task switching interference | 0.06 | Impulsiveness | 0.12 |
| Mania/ disorganization* | 0.15 | Reward dependence | 0.02 | Spatial WM load 7 accuracy* | 0.06 | Spatial manipulation accuracy* | 0.11 |
| Task switching interference | 0.15 | Positive formal thought disorder* | 0.01 | Delay discounting medium rewards* | 0.06 | Verbal maintenance RT* | 0.11 |
| Go/no go false alarm rate | 0.15 | Task switching residual switch cost | 0.01 | Mania* | 0.06 | Functional impulsivity | 0.10 |
| Attention* | 0.12 | Energy/ activity | 0.00 | Task switching switch cost | 0.06 | Verbal manipulation RT* | 0.10 |
| Spatial WM load 3 RT* | 0.11 | Remember RT* | -0.01 | Delay discounting total* | 0.05 | Hyperactivity* | 0.10 |
| Attention network task interference RT | 0.11 | Anhedonia* | -0.03 | Spatial manipulation accuracy* | 0.05 | Empathy | 0.08 |
| Scene recognition encoding RT* | 0.10 | Verbal WM load 7 RT* | -0.03 | Spatial WM load 3 accuracy* | 0.05 | Go/no go hits RT | 0.08 |
| Remember forced recognition 1 feature* | 0.10 | Stroop accuracy* | -0.05 | Verbal WM load 5 accuracy* | 0.04 | Verbal WM load 3 accuracy* | 0.07 |
| Verbal WM load 5 RT* | 0.10 | Know forced recognition 1 feature* | -0.06 | Spatial WM load 1 accuracy* | 0.04 | Spatial WM load 3 accuracy* | 0.06 |
| Task switching residual switch cost | 0.09 | Go/no go hits RT | -0.06 | Verbal WM load 7 RT* | 0.03 | Avolition* | 0.06 |
| Spatial WM load 5 RT* | 0.08 | Bizarre behavior* | -0.07 | Infrequency | 0.03 | Schizoid personality | 0.05 |
| Scene recognition recall RT* | 0.08 | Mania* | -0.07 | Obsessive compulsive | 0.03 | Dysfunctional impulsivity | 0.05 |
| Positive formal thought disorder* | 0.07 | Functional impulsivity | -0.07 | Symbol span | 0.03 | Attention* | 0.04 |
| Spatial maintenance RT* | 0.07 | Spanish verbal fluency* | -0.08 | Verbal WM max capacity* | 0.03 | Stop signal RT* | 0.04 |
| Go/no go hits RT | 0.06 | Verbal WM load 9 RT* | -0.08 | Spatial WM load 5 accuracy* | 0.02 | Symbol span | 0.04 |
| Hyperactivity* | 0.06 | Venturesomeness | -0.09 | Remember accuracy colors* | 0.02 | Go/no go false alarm rate | 0.03 |
| Verbal WM load 3 RT* | 0.06 | Stroop RT* | -0.09 | Perceptual aberrations | 0.02 | Spatial WM max capacity* | 0.03 |
| Spatial manipulation RT* | 0.05 | Spatial manipulation accuracy* | -0.11 | Digit span backward | 0.02 | Spatial WM load 5 accuracy* | 0.03 |
| Know forced recognition 1 feature* | 0.05 | Remember accuracy colors* | -0.11 | Delay discounting large rewards* | 0.01 | Spatial WM load 7 accuracy* | 0.02 |

| LC1 | | LC2 | | LC3 | | LC4 | |
|---|---|---|---|---|---|---|---|
| **Behavior** | **Corr** | **Behavior** | **Corr** | **Behavior** | **Corr** | **Behavior** | **Corr** |
| Spatial WM load 1 RT* | 0.05 | Know accuracy colors* | -0.11 | Visual reproduction delayed recall | 0.01 | Stroop accuracy* | 0.02 |
| Stop signal RT* | 0.04 | High risk pumps* | -0.14 | Know accuracy words* | 0.01 | ADHD symptoms | 0.02 |
| Stroop RT* | 0.03 | Spatial WM load 5 accuracy* | -0.14 | Anxiety | 0.01 | Control | 0.02 |
| Verbal maintenance RT* | 0.00 | Remember forced recognition 1 feature* | -0.14 | Verbal maintenance RT* | 0.01 | Spatial WM load 7 RT* | 0.01 |
| Spatial WM load 7 RT* | -0.01 | Anxiety | -0.16 | Daydreaming | 0.01 | Verbal WM load 5 accuracy* | 0.01 |
| Verbal WM load 7 RT* | -0.01 | Verbal WM load 9 accuracy* | -0.16 | Remember RT* | 0.01 | Verbal WM load 3 RT* | 0.01 |
| Task switching switch cost | -0.01 | Impulsiveness | -0.17 | Vocabulary | 0.01 | Spatial WM load 1 accuracy* | 0.01 |
| Color trail interference | -0.01 | Mania/ disorganization* | -0.17 | Stroop RT* | 0.01 | Scene recognition recall accuracy* | 0.01 |
| Delay discounting large rewards* | -0.02 | Verbal WM load 5 accuracy* | -0.17 | High risk pumps* | 0.00 | Alogia* | 0.01 |
| Delay discounting small rewards* | -0.03 | Somatization | -0.18 | Verbal WM load 3 RT* | 0.00 | Attentional impulsivity | 0.01 |
| Delay discounting total* | -0.03 | Spatial WM load 3 accuracy* | -0.18 | Scene recognition recall accuracy* | 0.00 | Motor impulsivity | 0.01 |
| Verbal manipulation RT* | -0.05 | Know forced recognition 2 features* | -0.18 | Verbal manipulation accuracy* | 0.00 | Know forced recognition 1 feature* | 0.00 |
| Remember RT* | -0.05 | Spatial WM load 1 accuracy* | -0.18 | Interpersonal sensitivity | 0.00 | Delay discounting large rewards* | 0.00 |
| Low risk pumps* | -0.05 | Empathy | -0.19 | Verbal WM load 5 RT* | -0.01 | Verbal WM load 9 RT* | 0.00 |
| Delay discounting medium rewards* | -0.05 | Hyperactivity* | -0.19 | Attention network task interference RT | -0.01 | Task switching switch cost | 0.00 |
| Total pumps* | -0.06 | Low risk pumps* | -0.19 | Attention* | -0.01 | Know RT* | -0.01 |
| Verbal WM load 9 RT* | -0.07 | Remember forced recognition 2 features* | -0.19 | Spatial WM max capacity* | -0.02 | Verbal WM load 5 RT* | -0.01 |
| Venturesomeness | -0.07 | Scene recognition encoding accuracy* | -0.20 | Task switching residual switch cost | -0.02 | Spatial WM load 1 RT* | -0.01 |
| Know RT* | -0.07 | Social anhedonia | -0.20 | Visual reproduction immediate recall | -0.02 | Spatial maintenance RT* | -0.01 |
| Know accuracy colors* | -0.08 | Total pumps* | -0.20 | Verbal WM load 9 accuracy* | -0.02 | Harm avoidance | -0.01 |
| Know accuracy words* | -0.08 | Know accuracy words* | -0.20 | Spatial WM load 7 RT* | -0.02 | Verbal WM load 7 RT* | -0.01 |
| High risk pumps* | -0.10 | Spatial WM max capacity* | -0.21 | Spatial WM load 3 RT* | -0.02 | Scene recognition encoding RT* | -0.01 |
| Spatial WM load 7 accuracy* | -0.12 | Spatial WM load 7 accuracy* | -0.21 | Total pumps* | -0.03 | Delay discounting total* | -0.02 |
| Remember accuracy words* | -0.12 | Verbal WM load 3 accuracy* | -0.22 | Scene recognition encoding accuracy* | -0.03 | Delay discounting small rewards* | -0.02 |
| Spatial WM max capacity* | -0.13 | Verbal maintenance accuracy* | -0.22 | Long delay recognition | -0.03 | Verbal WM load 7 accuracy* | -0.02 |
| Go/no go hit rate | -0.13 | Nonplanning | -0.22 | Mood lability | -0.03 | Inattention* | -0.03 |
| Verbal manipulation accuracy* | -0.13 | Hypomanic personality | -0.23 | Low risk pumps* | -0.03 | Know accuracy colors* | -0.03 |
| Functional impulsivity | -0.13 | Scene recognition recall accuracy* | -0.25 | Hallucinations* | -0.04 | Remember forced recognition 1 feature* | -0.03 |
| Know forced recognition 2 features* | -0.14 | Depression (HAMD)* | -0.25 | Verbal WM load 9 RT* | -0.04 | Delay discounting medium rewards* | -0.03 |

| LC1 | | LC2 | | LC3 | | LC4 | |
|---|---|---|---|---|---|---|---|
| **Behavior** | **Corr** | **Behavior** | **Corr** | **Behavior** | **Corr** | **Behavior** | **Corr** |
| Remember accuracy colors* | -0.14 | Dysfunctional impulsivity | -0.26 | Remember forced recognition 1 feature* | -0.04 | Long delay recognition | -0.03 |
| Spatial manipulation accuracy* | -0.14 | Harm avoidance | -0.26 | Spatial maintenance accuracy* | -0.04 | Spatial WM load 5 RT* | -0.04 |
| Digit span forward | -0.15 | Verbal WM load 7 accuracy* | -0.26 | Spatial WM load 1 RT* | -0.04 | Matrix reasoning | -0.04 |
| Verbal WM load 9 accuracy* | -0.16 | Depression/ anxiety* | -0.26 | Depression (HSCL) | -0.05 | Verbal maintenance accuracy* | -0.04 |
| Spatial WM load 1 accuracy* | -0.17 | Obsessive compulsive | -0.27 | Spatial manipulation RT* | -0.05 | Stroop RT* | -0.05 |
| Scene recognition encoding accuracy* | -0.18 | Spatial maintenance accuracy* | -0.27 | Remember forced recognition 2 features* | -0.05 | Verbal WM max capacity* | -0.05 |
| Remember forced recognition 2 features* | -0.18 | Verbal WM max capacity* | -0.28 | Delusions* | -0.05 | Scene recognition encoding accuracy* | -0.06 |
| Stroop accuracy* | -0.19 | Interpersonal sensitivity | -0.30 | Spatial WM load 5 RT* | -0.05 | Spatial manipulation RT* | -0.06 |
| Spatial WM load 3 accuracy* | -0.21 | Attentional impulsivity | -0.31 | Scene recognition recall RT* | -0.05 | Task switching residual switch cost | -0.06 |
| Spatial maintenance accuracy* | -0.21 | Remember accuracy words* | -0.31 | Depression (HAMD)* | -0.07 | Low risk pumps* | -0.07 |
| Verbal WM load 5 accuracy* | -0.21 | Depression (HSCL) | -0.32 | Short delay free recall | -0.07 | Know forced recognition 2 features* | -0.07 |
| Verbal WM max capacity* | -0.21 | Verbal manipulation accuracy* | -0.32 | Know RT* | -0.07 | Know accuracy words* | -0.07 |
| Vocabulary | -0.22 | Schizoid personality | -0.33 | Spatial maintenance RT* | -0.07 | Spatial WM load 3 RT* | -0.07 |
| English verbal fluency | -0.23 | Task switching accuracy | -0.33 | Know forced recognition 1 feature* | -0.08 | Color trail interference | -0.07 |
| Verbal WM load 7 accuracy* | -0.23 | Mood lability | -0.33 | Short delay cued recall | -0.08 | Total pumps* | -0.08 |
| Verbal WM load 3 accuracy* | -0.23 | Novelty seeking | -0.33 | Go/no go hit rate | -0.08 | Reward dependence | -0.09 |
| Spatial WM load 5 accuracy* | -0.23 | ADHD symptoms | -0.34 | Depression/ anxiety* | -0.08 | Scene recognition recall RT* | -0.10 |
| Digit span backward | -0.23 | Inattention* | -0.34 | Long delay cued recall | -0.09 | Task switching interference | -0.11 |
| Task switching accuracy | -0.25 | Motor impulsivity | -0.34 | Scene recognition encoding RT* | -0.10 | Remember accuracy colors* | -0.12 |
| Verbal maintenance accuracy* | -0.25 | Social anxiety | -0.35 | Somatization | -0.10 | High risk pumps* | -0.13 |
| Reward dependence | -0.27 | Visual reproduction recognition | -0.38 | Long delay free recall | -0.11 | Daydreaming | -0.13 |
| Matrix reasoning | -0.27 | Visual reproduction delayed recall | -0.42 | Stop signal RT* | -0.12 | Verbal manipulation accuracy* | -0.14 |
| Digit span sequencing | -0.27 | Letter/number sequencing | -0.43 | Remember accuracy words* | -0.12 | Novelty seeking | -0.14 |
| Spanish verbal fluency* | -0.29 | Digit span forward | -0.44 | Task switching accuracy | -0.13 | Verbal WM load 9 accuracy* | -0.14 |
| Scene recognition recall accuracy* | -0.29 | Daydreaming | -0.44 | Anhedonia* | -0.13 | Task switching accuracy | -0.15 |
| Persistence | -0.32 | Visual reproduction immediate recall | -0.48 | Blunt affect* | -0.14 | Venturesomeness | -0.16 |
| Visual reproduction recognition | -0.32 | Long delay recognition | -0.49 | Schizoid personality | -0.14 | Remember forced recognition 2 features* | -0.16 |

| LC1 | | LC2 | | LC3 | | LC4 | |
|---|---|---|---|---|---|---|---|
| **Behavior** | **Corr** | **Behavior** | **Corr** | **Behavior** | **Corr** | **Behavior** | **Corr** |
| Long delay recognition | -0.34 | Digit span sequencing | -0.49 | Alogia* | -0.19 | Digit span sequencing | -0.17 |
| Letter/number sequencing | -0.35 | Digit span backward | -0.50 | Positive symptoms* | -0.20 | Nonplanning | -0.19 |
| Symbol span | -0.39 | Symbol span | -0.51 | Avolition* | -0.24 | Remember accuracy words* | -0.19 |
| Visual reproduction delayed recall | -0.42 | English verbal fluency | -0.51 | Go/no go hits RT | -0.26 | Go/no go hit rate | -0.20 |
| Visual reproduction immediate recall | -0.43 | Short delay free recall | -0.52 | Physical anhedonia | -0.29 | English verbal fluency | -0.23 |
| Short delay cued recall | -0.49 | Long delay free recall | -0.54 | Negative symptoms* | -0.30 | Letter/number sequencing | -0.23 |
| Long delay free recall | -0.50 | Matrix reasoning | -0.55 | Social anhedonia | -0.31 | Attention network task interference RT | -0.30 |
| Long delay cued recall | -0.51 | Short delay cued recall | -0.56 | Social anxiety | -0.50 | Vocabulary | -0.32 |
| Short delay free recall | -0.52 | Long delay cued recall | -0.57 | Control | -0.55 | Digit span backward | -0.32 |
| Control | -0.60 | Vocabulary | -0.59 | Harm avoidance | -0.59 | Digit span forward | -0.40 |

*Abbreviations*: HAMD, Hamilton Psychiatric Rating Scale for Depression; HSCL, Hopkins Symptom Checklist; RT, reaction time; WM, working memory.

**Table S6. Absolute correlations between RSFC and behavioral composite scores obtained in reliability analyses**

| | Original PLS | Task fMRI PLS | | Cross-validated PLS | | |
|---|---|---|---|---|---|---|
| | | | | *Training data* | *Testing data* | |
| **Latent component** | **Corr** | **Corr** | **Permuted p** | **Corr (mean)** | **Corr (mean)** | **Permuted p (range)** |
| LC1 | 0.78 | 0.37 | 0.001 | 0.84 | 0.15 | 0.001 - 0.002 |
| LC2 | 0.83 | 0.40 | 0.001 | 0.84 | 0.12 | 0.002 - 0.004 |
| LC3 | 0.73 | 0.38 | 0.001 | 0.83 | 0.18 | 0.002 - 0.002 |
| LC4 | 0.80 | 0.42 | 0.001 | 0.85 | 0.14 | 0.001 - 0.002 |

(1) In the task fMRI PLS, task FC values were multiplied by the RSFC saliences (from the original PLS analysis) to obtain Task FC composite scores, which were then correlated with behavioral composite scores. (2) In the cross-validated PLS, we separated the data into a training (80% of the participants) and a test set (remaining 20% participants). PLS was first computed on the training data; then, the RSFC and behavior data of the test set were multiplied by the RSFC and behavioral saliences to obtain RSFC and behavioral composite scores for the test data, which were then correlated with each other. Statistical significance was assessed by permuting behavioral data 1000 times within each diagnostic group. For the cross-validated PLS analysis, correlations averaged across the 5 folds, as well as the range of p-values across folds are shown.

**Table S7. Absolute correlations between RSFC (or behavioral) saliences obtained in control analyses and RSFC (or behavioral) saliences from the original PLS analysis**

| | Latent component | *Behavior normalized* | *Skewed behaviors excluded* | *Confounds included* | *CompCor* | *Controls only* | *Patients only* |
|---|---|---|---|---|---|---|---|
| Correlations with original RSFC saliences | LC1 | 0.96 | 0.99 | 0.81 | 0.75 | 0.55 | 0.68 |
| | LC2 | 0.95 | 0.99 | 0.72 | 0.80 | 0.40 | 0.68 |
| | LC3 | 0.99 | 1.00 | 0.82 | 0.82 | 0.48 | 0.68 |
| | LC4 | 0.94 | 0.97 | 0.68 | 0.29 | 0.58 | 0.20 |
| Correlations with original behavioral saliences | LC1 | 0.98 | 1.00 | 0.97 | 0.83 | 0.76 | 0.97 |
| | LC2 | 0.93 | 0.99 | 0.79 | 0.91 | 0.80 | 0.89 |
| | LC3 | 0.98 | 1.00 | 0.90 | 0.96 | 0.76 | 0.92 |
| | LC4 | 0.97 | 0.98 | 0.75 | 0.56 | 0.80 | 0.22 |

These control analyses were performed after (1) quantile normalization was applied to all behavioral measures to improve Gaussianity ; (2) excluding four behavioral measures that had a skewed distribution ; (3) adding confounds (age, sex, years of education, acquisition site and head motion) to the rest of behavioral measures, instead of regressing them from the data prior to the PLS analysis ; (4) applying CompCor (16) instead of global signal regression (49) during rs-fMRI preprocessing ; (5) including healthy controls only ; (6) including patients only (all patient groups).

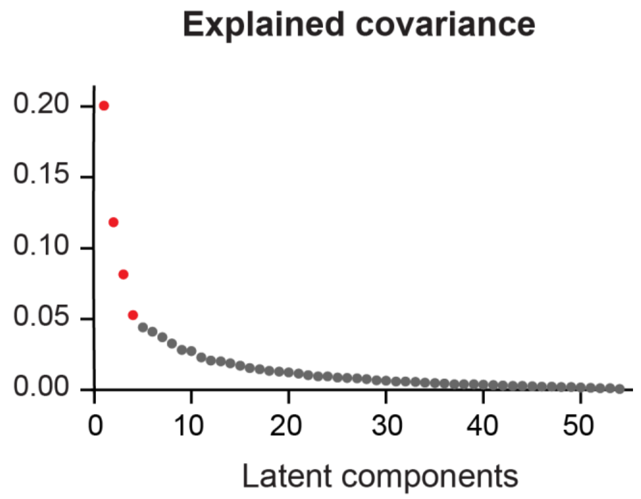


**Figure S1. Covariance explained by each latent component obtained with the PLS analysis.** The first four components (in red) were found to be statistically significant by permutation testing with FDR correction ($q < 0.05$).

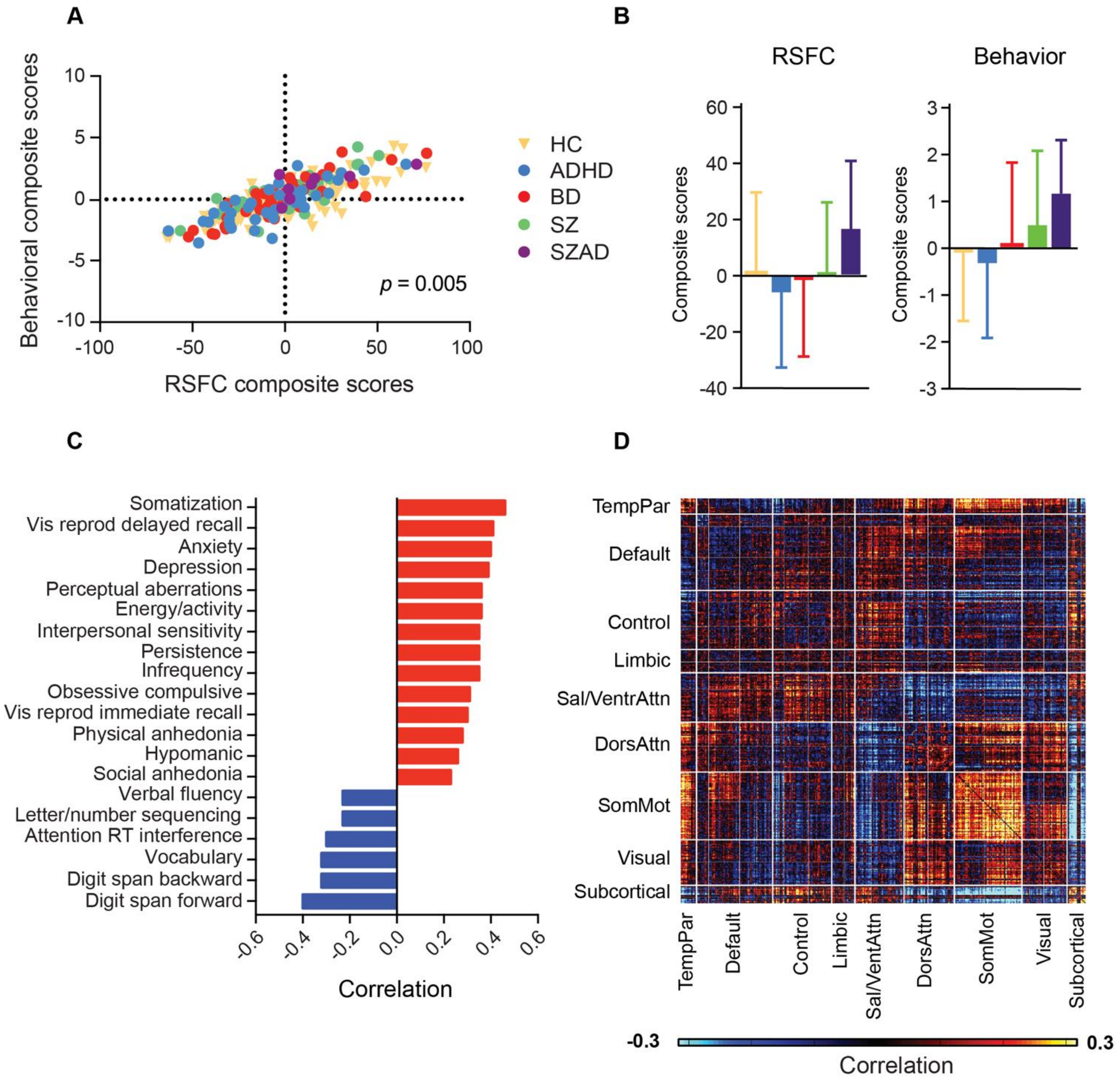


**Figure S2. Fourth latent component (LC4).** (A) Correlation between individual-specific RSFC and behavioral composite scores of participants. (B) Group differences in RSFC and behavioral composite scores. There was no significant group difference after FDR correction (q < 0.05). (C) Top 20 strongest correlations between participants' behavioral measures and their behavioral composite scores. Greater loading on LC4 was associated with higher measures of psychopathology and worse working memory and language abilities. (D) Correlations between participants' RSFC data and their RSFC composite scores. Red (or blue) color indicates that greater RSFC is positively (or negatively) associated with LC4. Greater RSFC composite score was associated with increased RSFC within the Somatomotor networks. Somatomotor networks also showed greater RSFC with the Temporo-Parietal network and Dorsal Attention B network. Moreover, Somatomotor A and Dorsal Attention networks also showed increased RSFC with Visual networks. Finally, subcortical regions showed increased FC with Control B, and decreased RSFC with Somatomotor networks.

*Abbreviations*: HC, healthy controls; ADHD, attention deficit/hyperactivity disorder; BD, bipolar disorder; SZ, schizophrenia; SZAD, schizoaffective disorder.

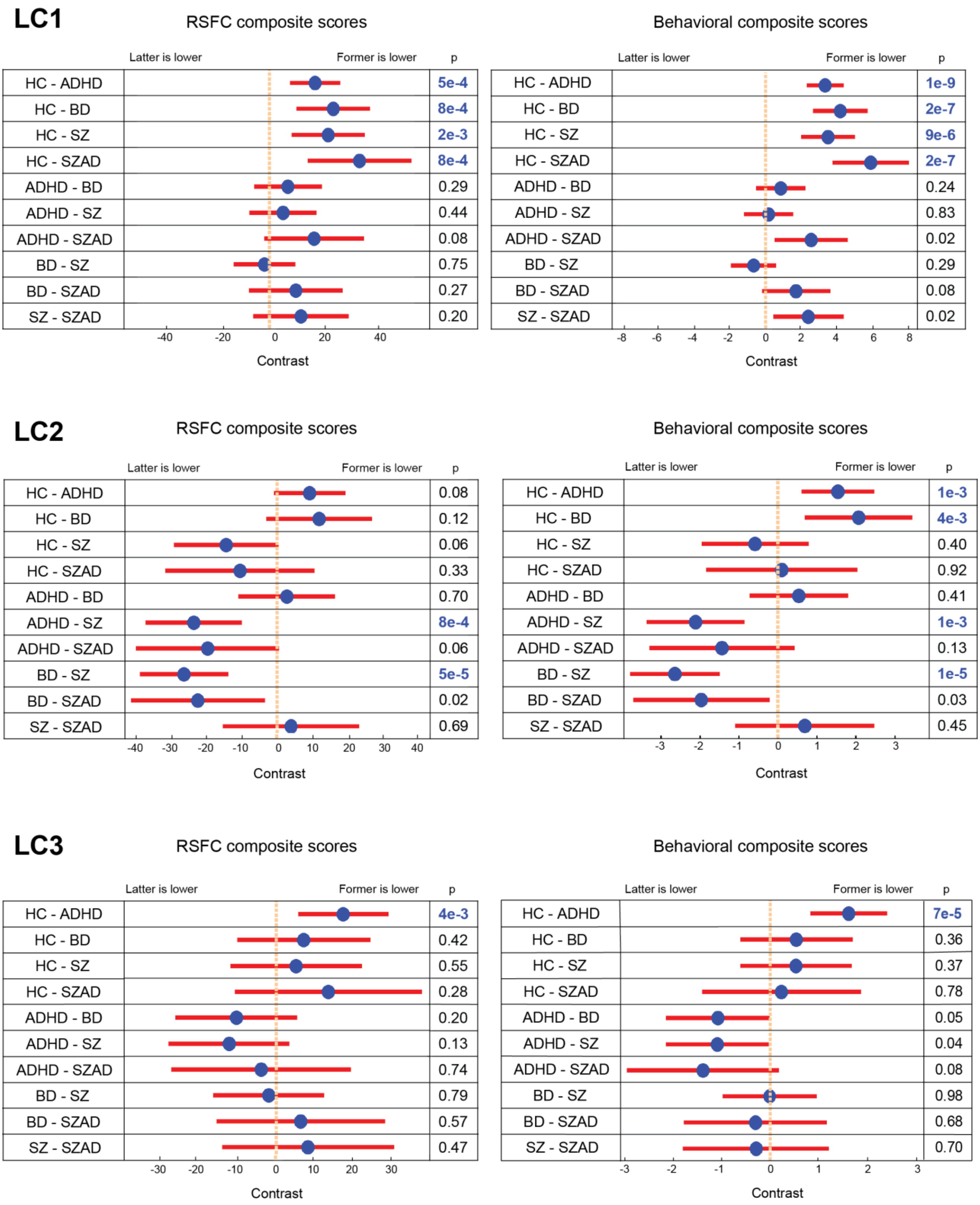


**Figure S3. Associations between diagnosis and RSFC (or behavioral) composite scores, while controlling for medication and substance use.** GLMs, followed by linear hypothesis tests, were used to compare betas coefficients of all pairs of diagnostic groups. All p-values that survived FDR correction ($q < 0.05$) are highlighted in blue. *Abbreviations*: HC, healthy controls; ADHD, attention deficit/hyperactivity disorder; BD, bipolar disorder; SZ, schizophrenia; SZAD, schizoaffective disorder.

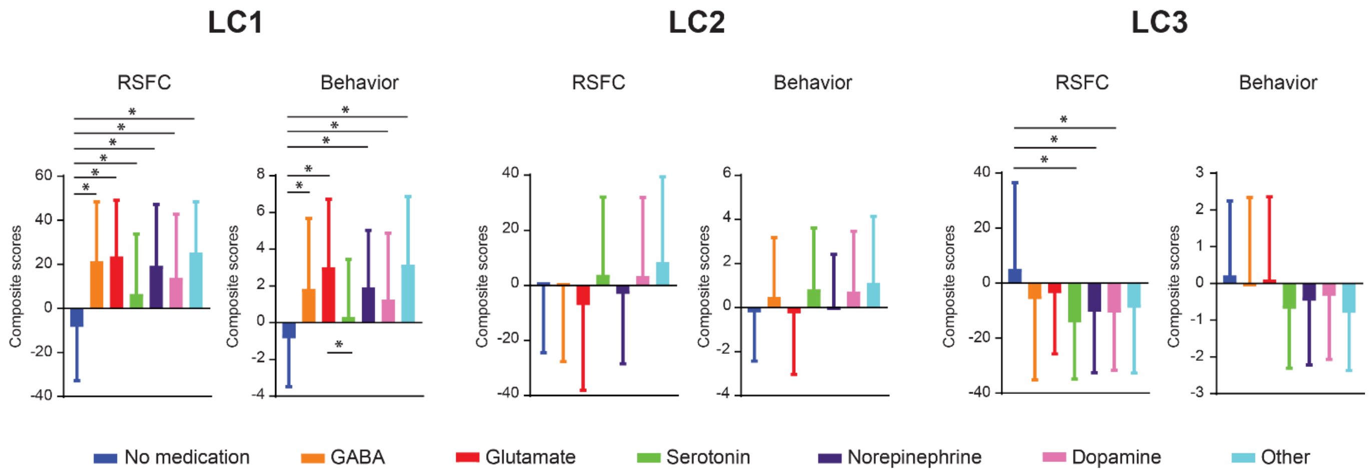


**Figure S4**. **Associations between medication (categorized by targeted neurotransmitter system) use and RSFC (or behavioral) composite scores.** T-tests were used to compare composite scores between all pairs of neurotransmitter systems targeted by the medication used by participants. All p-values that survived FDR correction ($q < 0.05$) are marked with *.

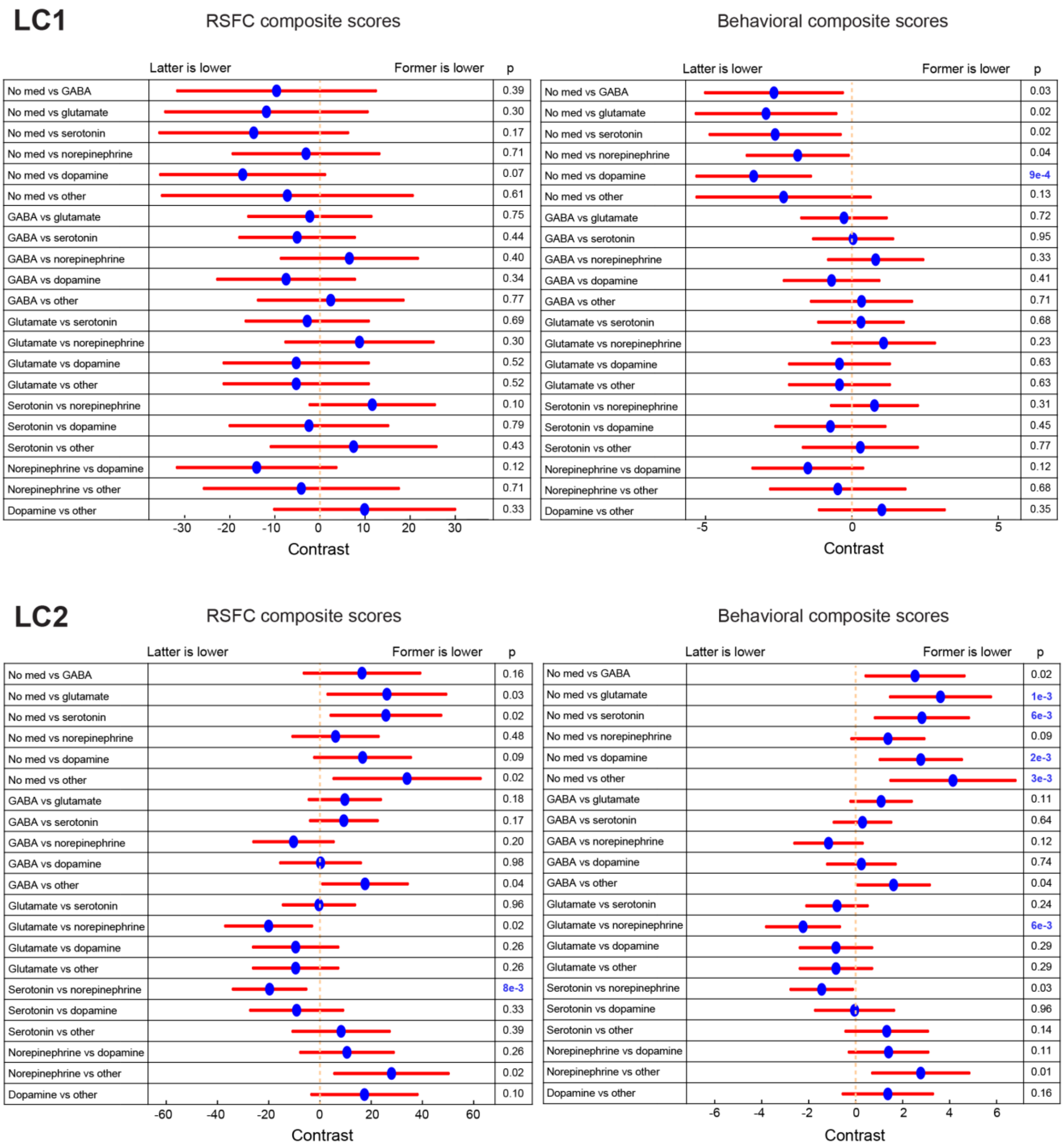
LC1
RSFC composite scores
Behavioral composite scores
Latter is lower
Former is lower
p
No med vs GABA 0.39 | 0.03
No med vs glutamate 0.30 | 0.02
No med vs serotonin 0.17 | 0.02
No med vs norepinephrine 0.71 | 0.04
No med vs dopamine 0.07 | 9e-4
No med vs other 0.61 | 0.13
GABA vs glutamate 0.75 | 0.72
GABA vs serotonin 0.44 | 0.95
GABA vs norepinephrine 0.40 | 0.33
GABA vs dopamine 0.34 | 0.41
GABA vs other 0.77 | 0.71
Glutamate vs serotonin 0.69 | 0.68
Glutamate vs norepinephrine 0.30 | 0.23
Glutamate vs dopamine 0.52 | 0.63
Glutamate vs other 0.52 | 0.63
Serotonin vs norepinephrine 0.10 | 0.31
Serotonin vs dopamine 0.79 | 0.45
Serotonin vs other 0.43 | 0.77
Norepinephrine vs dopamine 0.12 | 0.12
Norepinephrine vs other 0.71 | 0.68
Dopamine vs other 0.33 | 0.35
-30 -20 -10 0 10 20 30
-5 0 5
Contrast
LC2
RSFC composite scores
Behavioral composite scores
Latter is lower
Former is lower
p
No med vs GABA 0.16 | 0.02
No med vs glutamate 0.03 | 1e-3
No med vs serotonin 0.02 | 6e-3
No med vs norepinephrine 0.48 | 0.09
No med vs dopamine 0.09 | 2e-3
No med vs other 0.02 | 3e-3
GABA vs glutamate 0.18 | 0.11
GABA vs serotonin 0.17 | 0.64
GABA vs norepinephrine 0.20 | 0.12
GABA vs dopamine 0.98 | 0.74
GABA vs other 0.04 | 0.04
Glutamate vs serotonin 0.96 | 0.24
Glutamate vs norepinephrine 0.02 | 6e-3
Glutamate vs dopamine 0.26 | 0.29
Glutamate vs other 0.26 | 0.29
Serotonin vs norepinephrine 8e-3 | 0.03
Serotonin vs dopamine 0.33 | 0.96
Serotonin vs other 0.39 | 0.14
Norepinephrine vs dopamine 0.26 | 0.11
Norepinephrine vs other 0.02 | 0.01
Dopamine vs other 0.10 | 0.16
-60 -40 -20 0 20 40 60
-6 -4 -2 0 2 4 6
Contrast

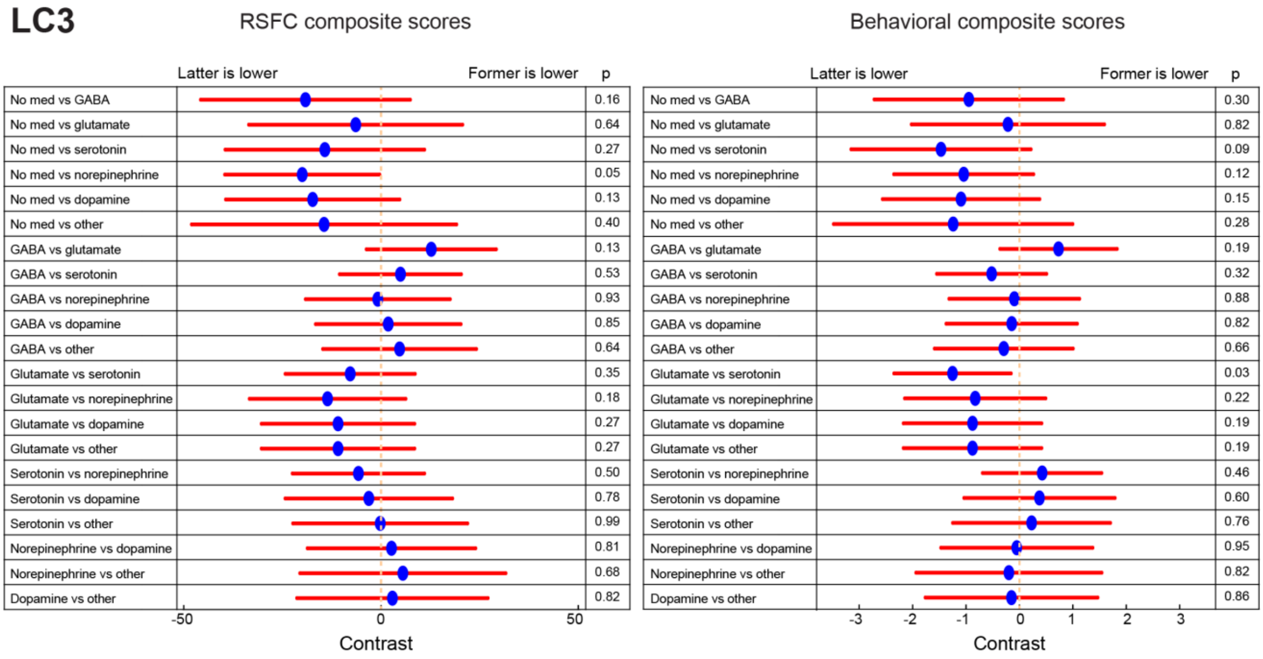


**Figure S5**. **Associations between medication (categorized by targeted neurotransmitter system) use and RSFC (or behavioral) composite scores, while controlling for diagnosis and substance use.** GLMs, followed by linear hypothesis tests, were used to compare betas coefficients of all pairs of neurotransmitter systems targeted by the medication used by participants. All p-values that survived FDR correction (q < 0.05) are highlighted in blue.

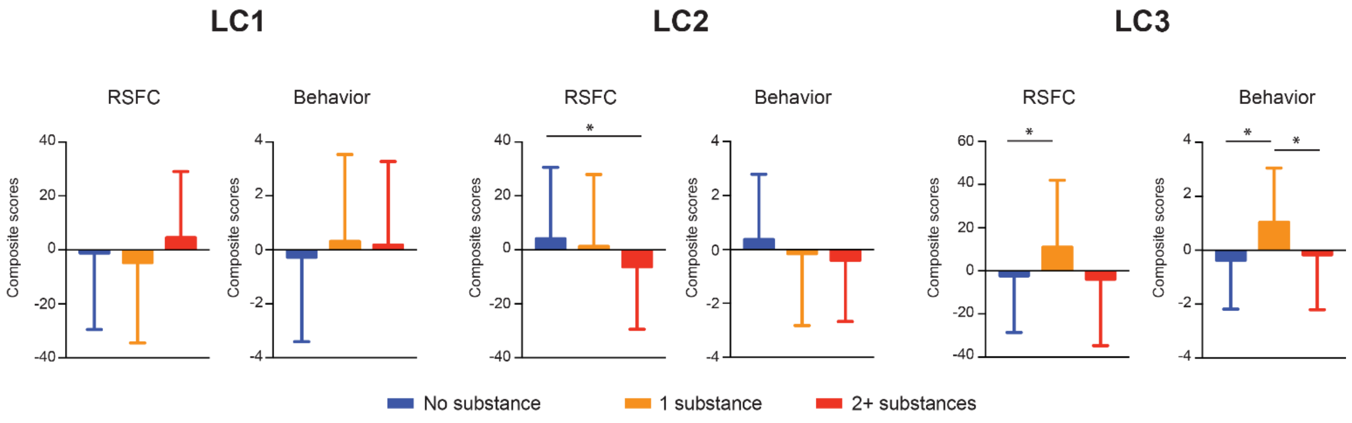


**Figure S6**. **Associations between substance use and RSFC (or behavioral) composite scores**. T-tests were used to compare composite scores between all pairs of substance categories (lifetime substance abuse and/or dependence on 0/1/2+ substances). Substances included nicotine, alcohol, cannabis, cocaine, amphetamine, sedatives / hypnotics / anxiolytics, inhalants, opioids and hallucinogens. All p-values that survived FDR correction ($q < 0.05$) are marked with *.

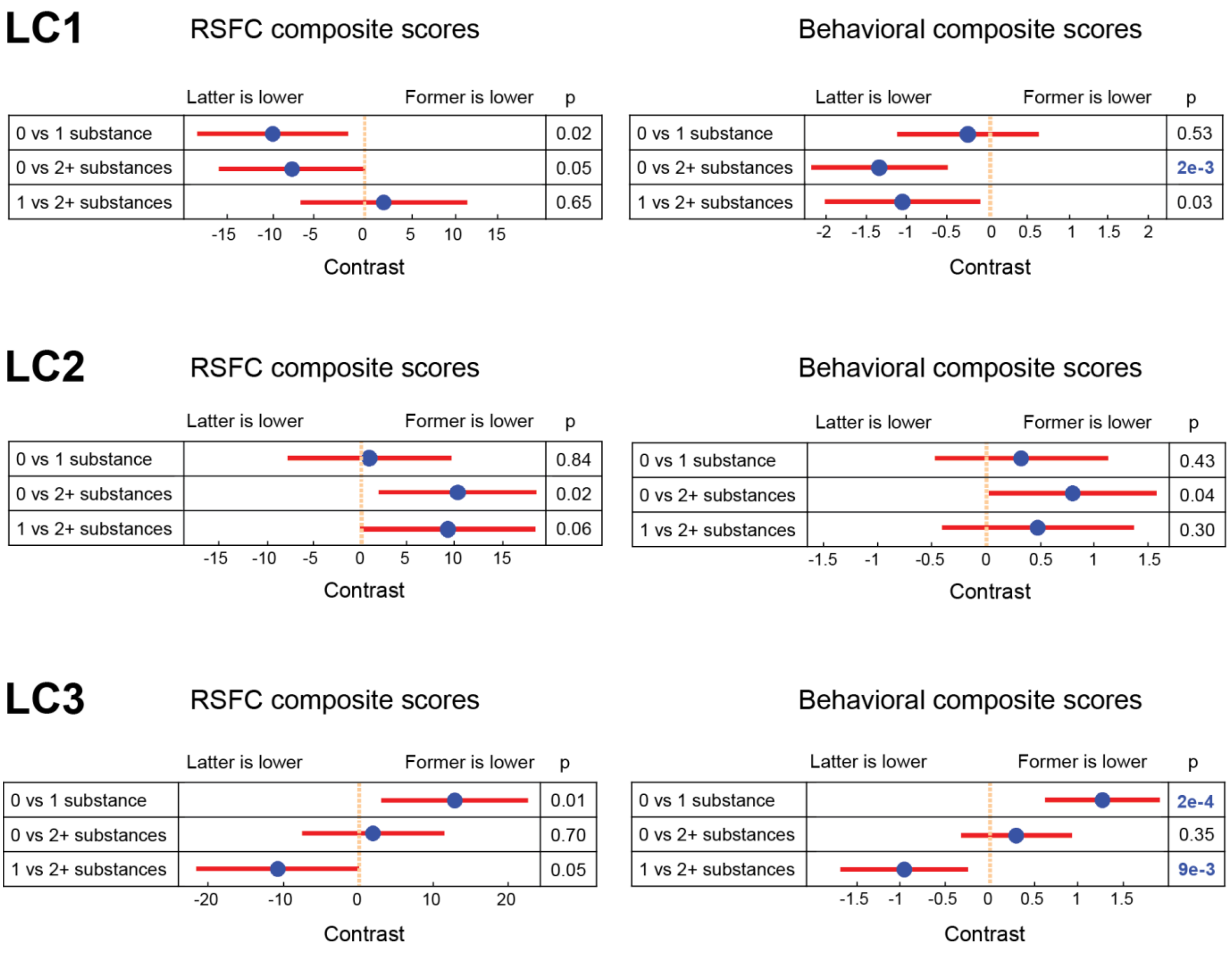


**Figure S7**. **Associations between substance use and RSFC (or behavioral) composite scores, while controlling for diagnosis and medication use**. GLMs, followed by linear hypothesis tests, were used to compare betas coefficients of all pairs of substance categories (lifetime abuse and/or dependence in 0/1/2+ substances). Substances included nicotine, alcohol, cannabis, cocaine, amphetamine, sedatives / hypnotics / anxiolytics, inhalants, opioids and hallucinogens. All p-values that survived FDR correction ($q < 0.05$) are highlighted in blue.

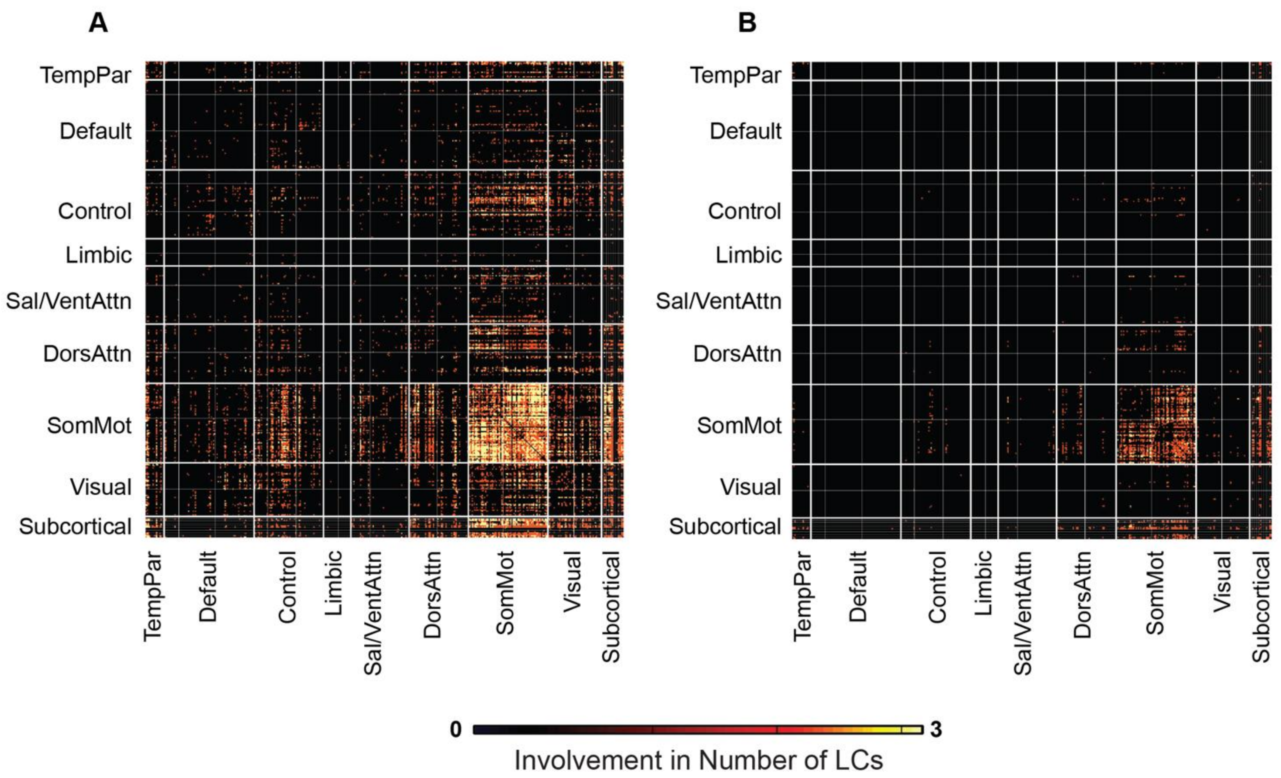


**Figure S8**. **Conjunction map showing common RSFC patterns across of LCs 1-2-3.** Absolute correlations between RSFC data and RSFC composite scores of LCs 1-2-3 (**Figures 1D, 2D and 3D**) were thresholded at 0.2 (A) and 0.3 (B). Connections that survived the threshold in only one map were changed to 0 (black); those that survived the threshold in two or three maps had a value of 2 (orange) and 3 (yellow), respectively.

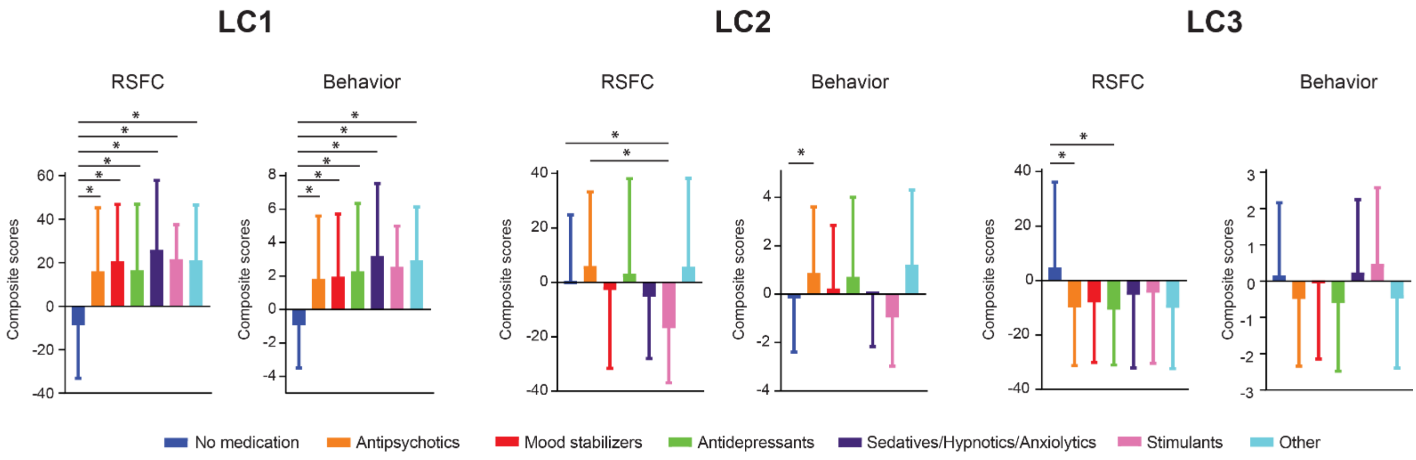


**Figure S9**. **Associations between medication (categorized by medication class) use and RSFC (or behavioral) composite scores.** T-tests were used to compare composite scores between all pairs of medication classes. All p-values that survived FDR correction (q < 0.05) are marked with *.

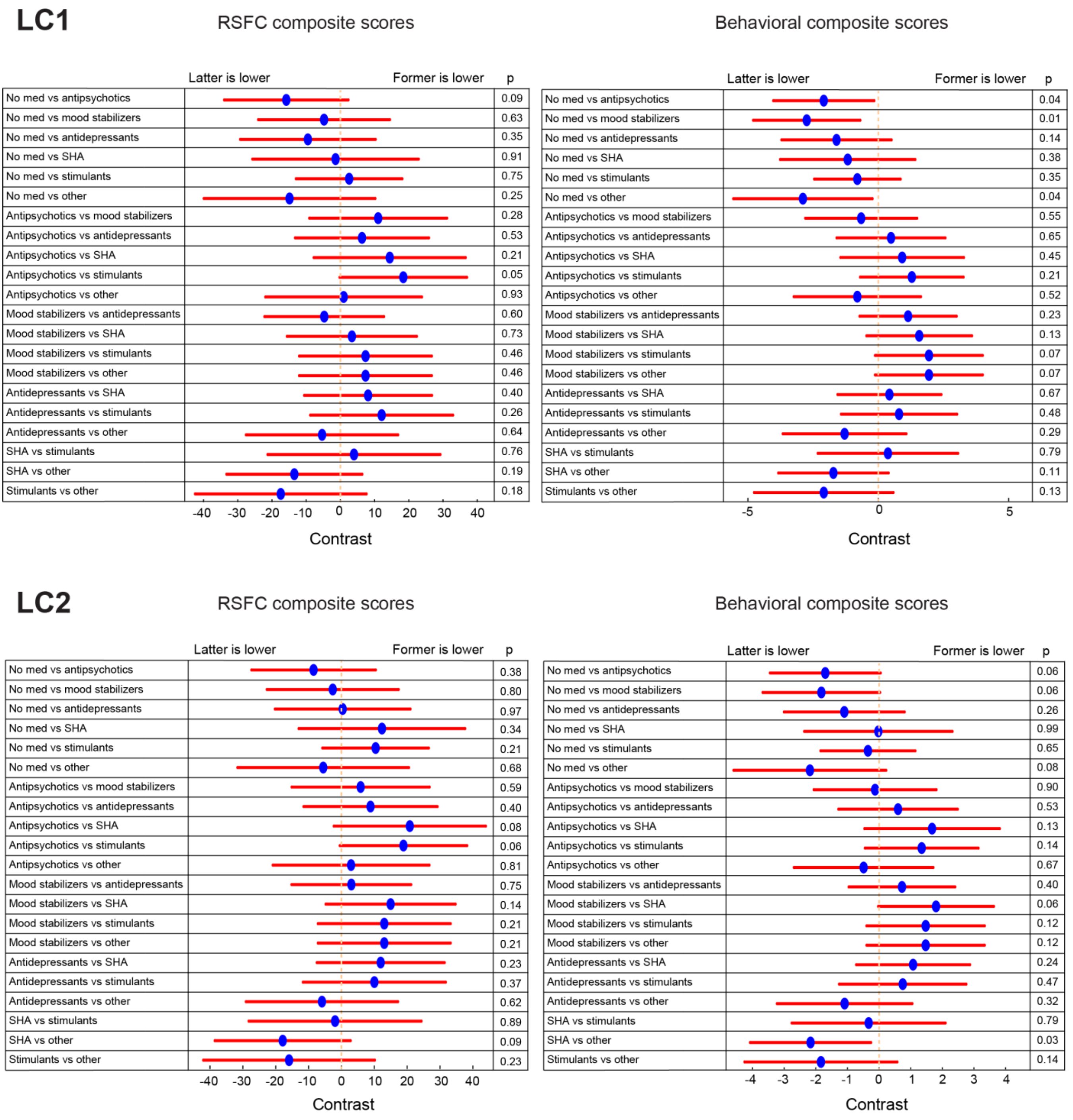
LC1
RSFC composite scores
Behavioral composite scores
Latter is lower
Former is lower
p
No med vs antipsychotics 0.09 | 0.04
No med vs mood stabilizers 0.63 | 0.01
No med vs antidepressants 0.35 | 0.14
No med vs SHA 0.91 | 0.38
No med vs stimulants 0.75 | 0.35
No med vs other 0.25 | 0.04
Antipsychotics vs mood stabilizers 0.28 | 0.55
Antipsychotics vs antidepressants 0.53 | 0.65
Antipsychotics vs SHA 0.21 | 0.45
Antipsychotics vs stimulants 0.05 | 0.21
Antipsychotics vs other 0.93 | 0.52
Mood stabilizers vs antidepressants 0.60 | 0.23
Mood stabilizers vs SHA 0.73 | 0.13
Mood stabilizers vs stimulants 0.46 | 0.07
Mood stabilizers vs other 0.46 | 0.07
Antidepressants vs SHA 0.40 | 0.67
Antidepressants vs stimulants 0.26 | 0.48
Antidepressants vs other 0.64 | 0.29
SHA vs stimulants 0.76 | 0.79
SHA vs other 0.19 | 0.11
Stimulants vs other 0.18 | 0.13
-40 -30 -20 -10 0 10 20 30 40
-5 0 5
Contrast
LC2
RSFC composite scores
Behavioral composite scores
Latter is lower
Former is lower
p
No med vs antipsychotics 0.38 | 0.06
No med vs mood stabilizers 0.80 | 0.06
No med vs antidepressants 0.97 | 0.26
No med vs SHA 0.34 | 0.99
No med vs stimulants 0.21 | 0.65
No med vs other 0.68 | 0.08
Antipsychotics vs mood stabilizers 0.59 | 0.90
Antipsychotics vs antidepressants 0.40 | 0.53
Antipsychotics vs SHA 0.08 | 0.13
Antipsychotics vs stimulants 0.06 | 0.14
Antipsychotics vs other 0.81 | 0.67
Mood stabilizers vs antidepressants 0.75 | 0.40
Mood stabilizers vs SHA 0.14 | 0.06
Mood stabilizers vs stimulants 0.21 | 0.12
Mood stabilizers vs other 0.21 | 0.12
Antidepressants vs SHA 0.23 | 0.24
Antidepressants vs stimulants 0.37 | 0.47
Antidepressants vs other 0.62 | 0.32
SHA vs stimulants 0.89 | 0.79
SHA vs other 0.09 | 0.03
Stimulants vs other 0.23 | 0.14
-40 -30 -20 -10 0 10 20 30 40
-4 -3 -2 -1 0 1 2 3 4
Contrast

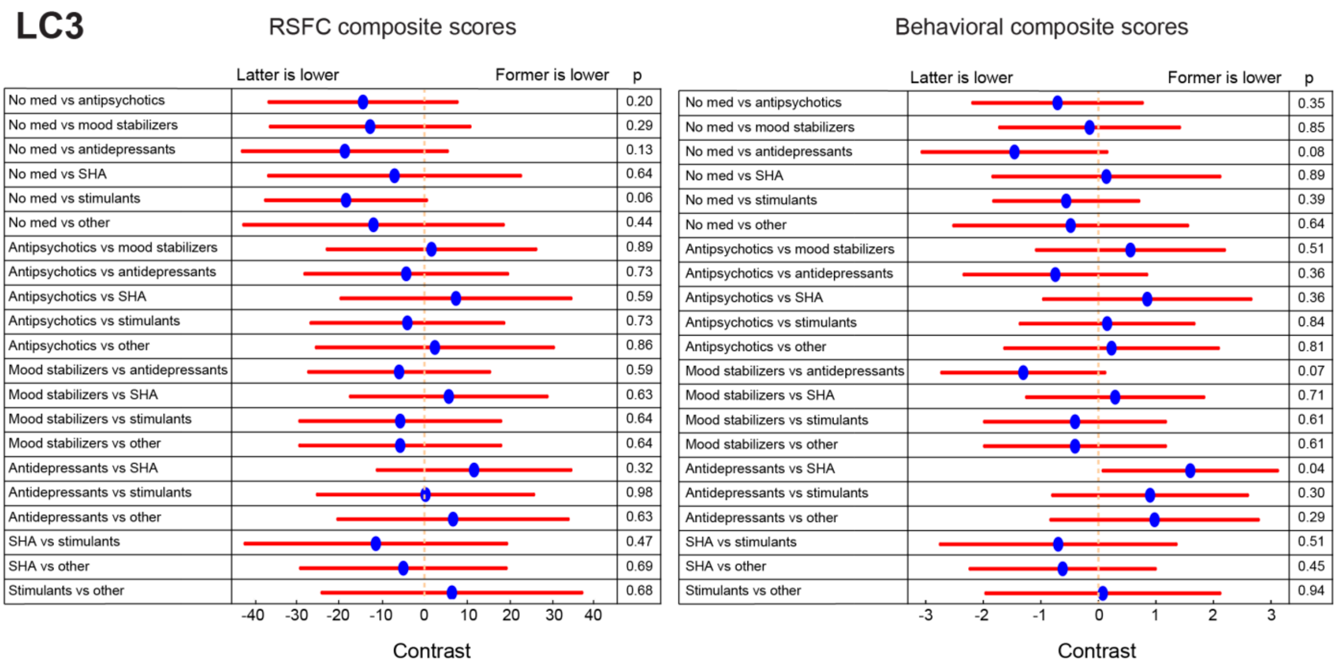


**Figure S10**. **Associations between medication (categorized by medication class) use and RSFC (or behavioral) composite scores, while controlling for diagnosis and substance use**. GLMs, followed by linear hypothesis tests, were used to compare betas coefficients of all pairs of medication classes used by participants. None of the p-values survived FDR correction (q < 0.05). *Abbreviations*: SHA, sedatives/hypnotics/anxiolytics.

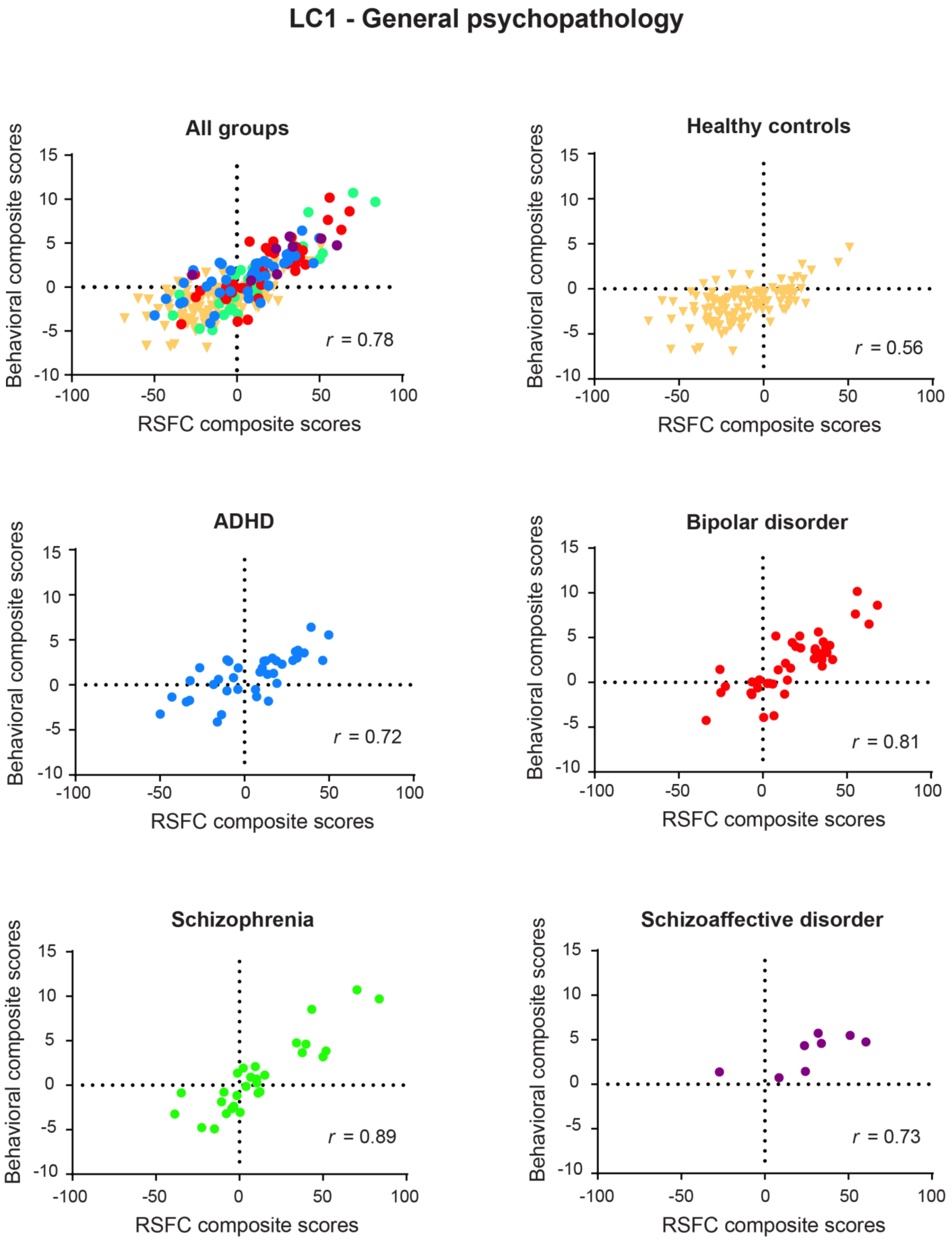


**Figure S11**. **Correlation between individual-specific RSFC and behavioral composite scores of the first latent component in each primary diagnostic group**

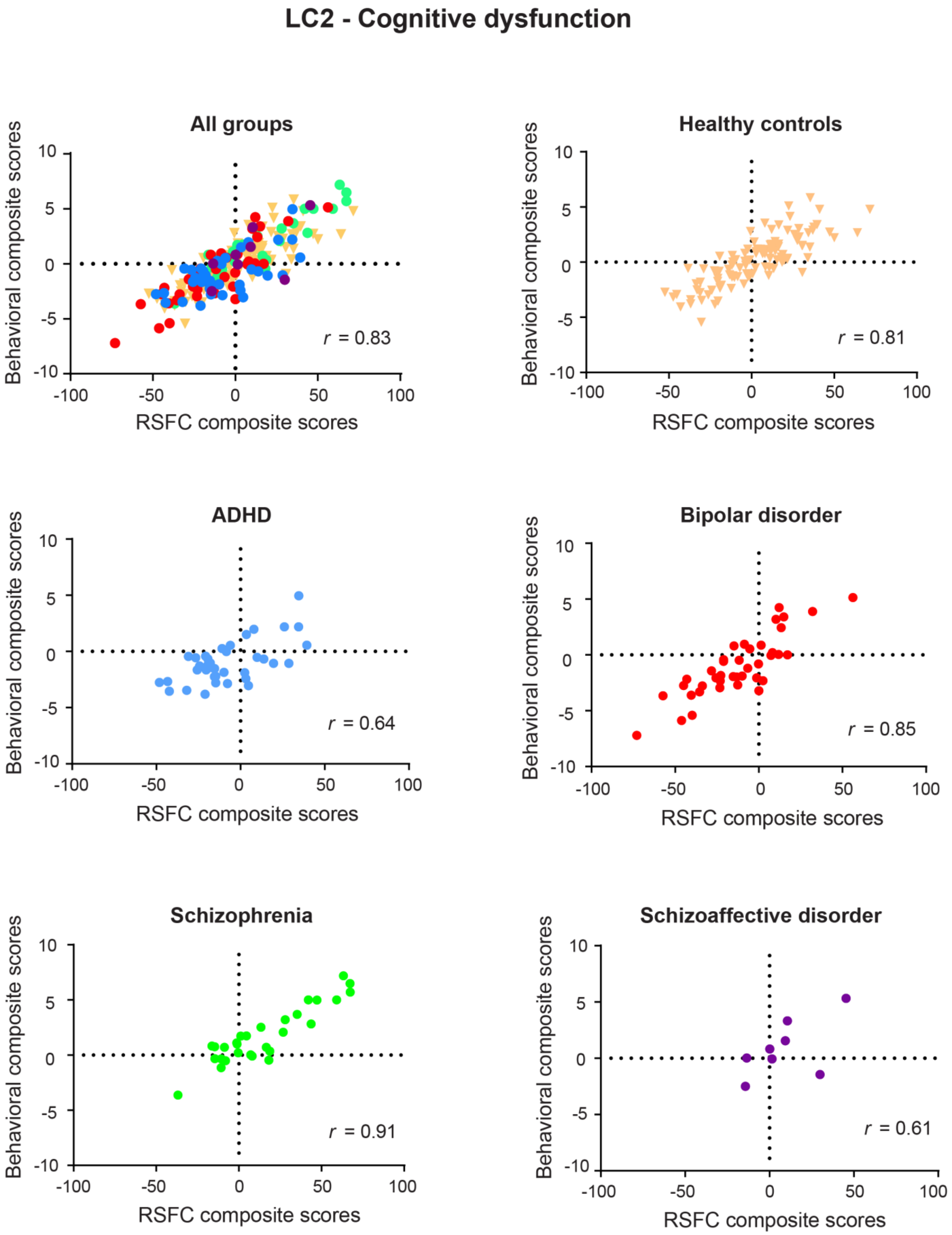


**Figure S12**. **Correlation between individual-specific RSFC and behavioral composite scores of the second latent component in each primary diagnostic group**

### LC3 - Impulsivity

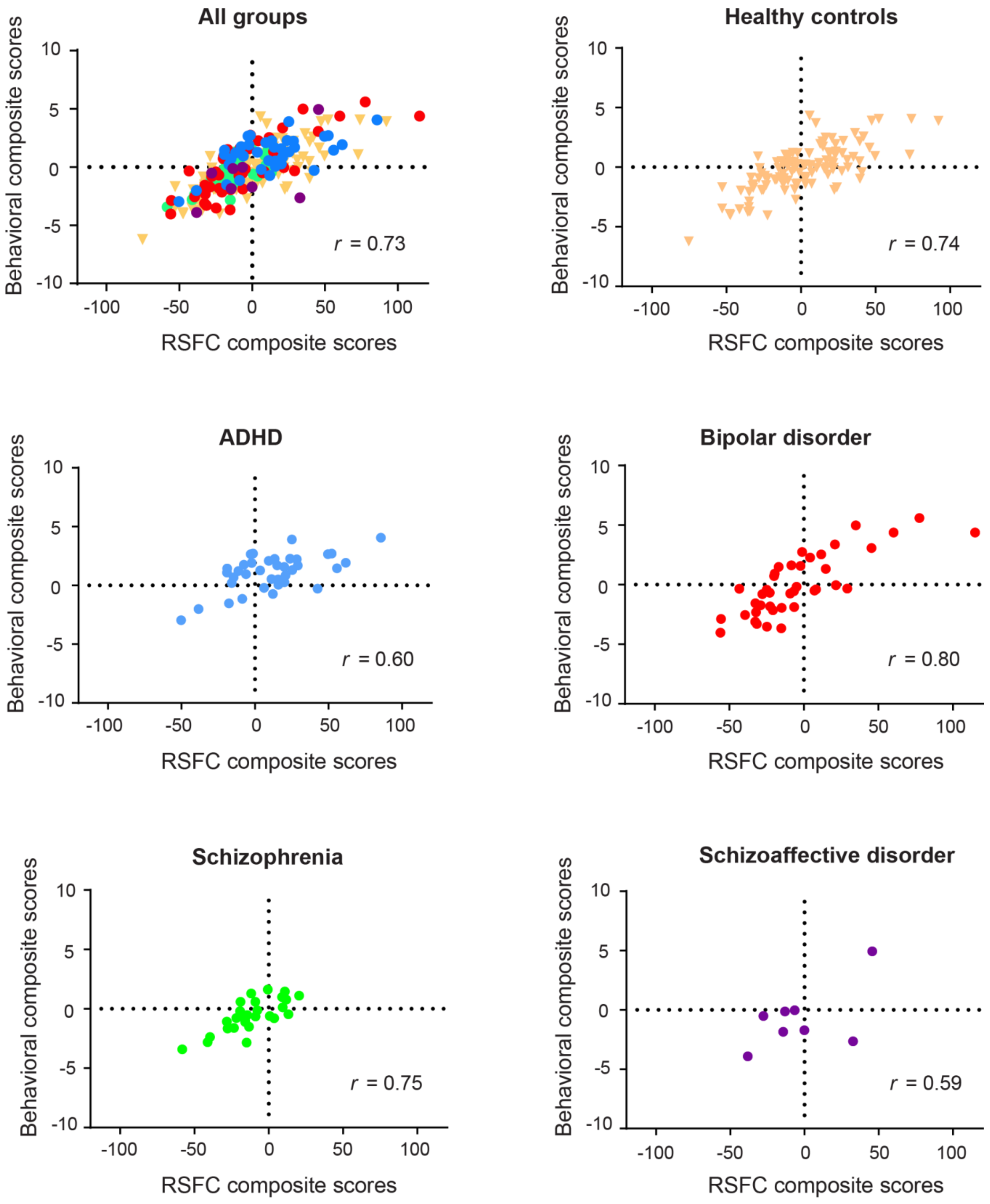


**Figure S13. Correlation between individual-specific RSFC and behavioral composite scores of the third latent component in each primary diagnostic group**
